# Two-dimensional perisaccadic visual mislocalization in rhesus macaque monkeys

**DOI:** 10.1101/2024.11.20.624548

**Authors:** Matthias P. Baumann, Ziad M. Hafed

**Affiliations:** Werner Reichardt Centre for Integrative Neuroscience, University of Tübingen, Tübingen, Germany, 72076; Hertie Institute for Clinical Brain Research, University of Tübingen, Tübingen, Germany, 72076

## Abstract

Perceptual localization of brief, high contrast perisaccadic visual probes is grossly erroneous. While this phenomenon has been extensively studied in humans, more needs to be learned about its underlying neural mechanisms. This ideally requires running similar behavioral paradigms in animals. However, during neurophysiology, neurons encountered in the relevant sensory and sensory-motor brain areas for visual mislocalization can have arbitrary, non-cardinal response field locations. This necessitates using mislocalization paradigms that can work with any saccade direction. Here, we first established such a paradigm in three male rhesus macaque monkeys. In every trial, the monkeys generated a visually-guided saccade towards an eccentric target. Once a saccade onset was detected, we presented a brief flash at one of three possible locations ahead of the saccade target location. After an experimentally-imposed delay period, we removed the saccade target, and the monkeys were then required to generate a memory-guided saccade towards the remembered flash location. All three monkeys readily learned the task, and, like humans, they all showed strong backward mislocalization towards the saccade target, which recovered for later flashes from the saccade time. Importantly, we then replicated a well-known property of human perisaccadic mislocalization, as revealed by two-dimensional mislocalization paradigms: that mislocalization is strongest for upward saccades. For horizontal saccades, we additionally found stronger mislocalization for upper visual field flashes, again consistent with humans. Our results establish a robust two-dimensional mislocalization paradigm in monkeys, and they pave the way for exploring the neural mechanisms underlying the dependence of perisaccadic mislocalization strength on saccade direction.

**Significance statement:** Visual perception is strongly altered around the time of saccades. Such alteration is often studied by characterizing how brief perisaccadic visual flashes are perceptually localized. While the properties of visual mislocalization have been exhaustively studied, the underlying neural mechanisms remain elusive, and this is due to a lack of suitable behavioral paradigms in animal models. We describe such a paradigm for macaques, which are ideal for exploring sensory-motor neural processes related to perisaccadic mislocalization.

## Introduction

Studying visual perception requires investigating the interplays between visual sensory processing and eye movements. This is because eye movements, such as saccades, can drastically alter retinal images even if the external environment is completely stable.

Multiple alterations in visual information processing are known to accompany saccades. For example, perisaccadic visual sensitivity is reduced, sometimes rendering brief visual presentations completely invisible (Latour, 1962; Zuber and Stark, 1966; Beeler, 1967). Moreover, when stimulus contrasts are high enough to be detected, these stimuli are grossly mislocalized (Honda, 1989; Cai et al., 1997; Ross et al., 1997). Such mislocalization depends on various factors related to the visual environment (Lappe et al., 2000). For example, in complete darkness, brief presaccadic flashes are perceived to have appeared ahead of their true positions (Dassonville et al., 1992; Lappe et al., 2000); such mislocalization is often referred to as forward mislocalization, since the error in reported position has a vector of similar direction to the saccade vector direction (Klingenhoefer and Krekelberg, 2017). Interestingly, the mislocalization is biphasic (Honda, 1991, 1993), with postsaccadic flashes being perceived as having been shifted opposite, rather than along, the saccade vector. On the other hand, with visual references, perisaccadic mislocalization is no longer biphasic, and appears like a “compression” towards the saccade target: brief flashes presented ahead of the saccade target experience mislocalization in the opposite direction from the saccade, even presaccadically (Ross et al., 1997; Klingenhoefer and Krekelberg, 2017); flashes presented in between the initial gaze position and the saccade target undergo forward mislocalization. Such “compression” could reflect the importance of the saccade goal in updating visual-motor reference frames (Schoppik and Lisberger, 2006; Zirnsak et al., 2014), and it implies that in structured visual environments, observing forward or backward mislocalization is no longer a function of time (like in complete darkness), but a function of spatial location (Honda, 1999).

Even though much of the early work on perisaccadic mislocalization has focused on only one dimension (horizontal saccades and perceptual reports on only the horizontal position of flashes), subsequent pioneering work demonstrated that, even with horizontal saccades, mislocalization has a distinct two-dimensional landscape (Kaiser and Lappe, 2004). More importantly, saccade directions themselves alter the two-dimensional mislocalization landscape, with upward saccades causing particularly strong reverse mislocalization for flashes presented ahead of the saccade target (Grujic et al., 2018). This latter result, which was motivated by a difference between upper and lower visual field representations in the superior colliculus (Hafed and Chen, 2016; Zhang et al., 2022; Hafed, 2025), suggests a clear need to study perisaccadic mislocalization, whether perceptually or with neurophysiological experiments, using more than just horizontal saccades.

Here, we aimed to establish a robust and flexible perisaccadic visual mislocalization paradigm in macaque monkeys, setting the stage for subsequent neurophysiological experiments. Four previous studies demonstrated perisaccadic mislocalization in monkeys, but all used only horizontal saccades and with only one-dimensional mislocalization measures (analyzing the horizontal component of flash positions). In one study, forward mislocalization was observed in complete darkness (Dassonville et al., 1992), consistent with human studies (Honda, 1989; Lappe et al., 2000). In another, backward mislocalization was observed in the absence of complete darkness (even for flashes behind the saccade target location), inconsistent with so-called “compression” (Jeffries et al., 2007). However, the flashes used in that study were not the very brief flashes typical of mislocalization experiments in humans. In the third study, perisaccadic “compression” was observed like in humans, and under identical experimental conditions between species (Klingenhoefer and Krekelberg, 2017). And, most recently, evidence of “compression” was again documented, enabling simultaneous recordings of visual cortical neurons during the behavior (Weng et al., 2024). Thus, it is indeed plausible to use monkeys for mislocalization experiments.

Having said that, the reality of neurophysiological experiments is that saccade direction could be completely arbitrary. For example, if one were to record superior colliculus saccade motor bursts, as a potential source of corollary discharge signals (Sommer and Wurtz, 2004, 2006; Berman et al., 2009; Berman and Wurtz, 2010, 2011; Berman et al., 2017) that may contribute to mislocalization, then it is more likely to record from oblique rather than horizontal saccade representations. This is also true when probing the representations of brief perisaccadic flashes themselves. Therefore, and given the dependence of perisaccadic mislocalization strength on saccade direction (Grujic et al., 2018), true translation of the classic human mislocalization studies to neurophysiological experiments would only be possible with a general, and robust, two-dimensional mislocalization paradigm in monkeys. This is what we document here.

## Materials and Methods

### Experimental animals and ethical approvals

We collected data from three adult, male rhesus macaque monkeys (*Macaca mulatta*). The monkeys (A, F, and M) were aged 11-15 years, and they weighed 10-13 kg.

All experiments were approved by the ethics committees at the regional governmental offices of the city of Tübingen.

### Laboratory setup and animal procedures

The monkeys were seated in front of a brightly-lit computer-controlled display, spanning approximately 31 deg horizontally and 23 deg vertically. The display was at a distance of ∼72 cm from the animals. We tracked eye movements with high precision using the scleral search coil technique (Robinson, 1963; Fuchs and Robinson, 1966; Judge et al., 1980), and the experiment was controlled in real-time using a modified version of PLDAPS (Eastman and Huk, 2012). Stimulus presentation and control were performed via the Psychophysics Toolbox (Brainard, 1997; Pelli, 1997; Kleiner et al., 2007), which provided exact time stamps at every frame update, ensuring accurate synchronization with the refresh cycle of the display. Prior tests with photodiodes (during the first few years of existence of the Hafed laboratory) confirmed the accurate timings of the Psychophysics Toolbox time, as expected given that the toolbox directly accesses the vertical blanking signal of the graphics card.

The animals were prepared for behavioral and neurophysiological experiments earlier (Tian et al., 2018; Buonocore et al., 2019; Skinner et al., 2019). This entailed a first surgery to implant a head fixation apparatus. During data collection, this apparatus was attached to a coupled device on the monkey chair in order to stabilize head position. Then, a later surgery was performed to implant a scleral search coil in one eye for eye movement tracking purposes.

### Experimental procedures

The main task involved generating a primary visually-guided saccade, around which a very brief probe flash was presented; later on in the very same trial, the monkeys generated a “report” saccade aiming at where they remembered the perisaccadic flash to have appeared.

Each trial (Fig. 1) started with an initial white fixation spot (of 79.9 cd/m^2^ luminance) presented over a uniform gray background (26.11 cd/m^2^). This illumination created a clear border along the edges of the display, providing a consistent visual reference frame throughout the experiments; thus, we were measuring “compression”-style mislocalization (Ross et al., 1997; Lappe et al., 2000). The fixation spot was displaced from display center (by 7 deg, except for the cardinal saccade directions of monkey M, in which case it was by 5 deg), and it could appear in one of eight (equally spaced) directions from this center. After 600-1000 ms, the fixation spot disappeared, and a simultaneous saccade target appeared at the display center. This target consisted of a vertical Gabor grating of either 0.5 cycles/deg (cpd) or 8 cycles/deg (cpd) spatial frequency, and it had a radius of 1.5 deg. The grating had high contrast (100%), and it featured a white central marker surrounded by a small gray disc, as in our previous experiments (Baumann et al., 2023). This central marker aided in maintaining saccade accuracy and precision across trials having different image appearances of the saccade target. The onset of the grating was the cue to generate a visually-guided saccade.

**Figure 1.**
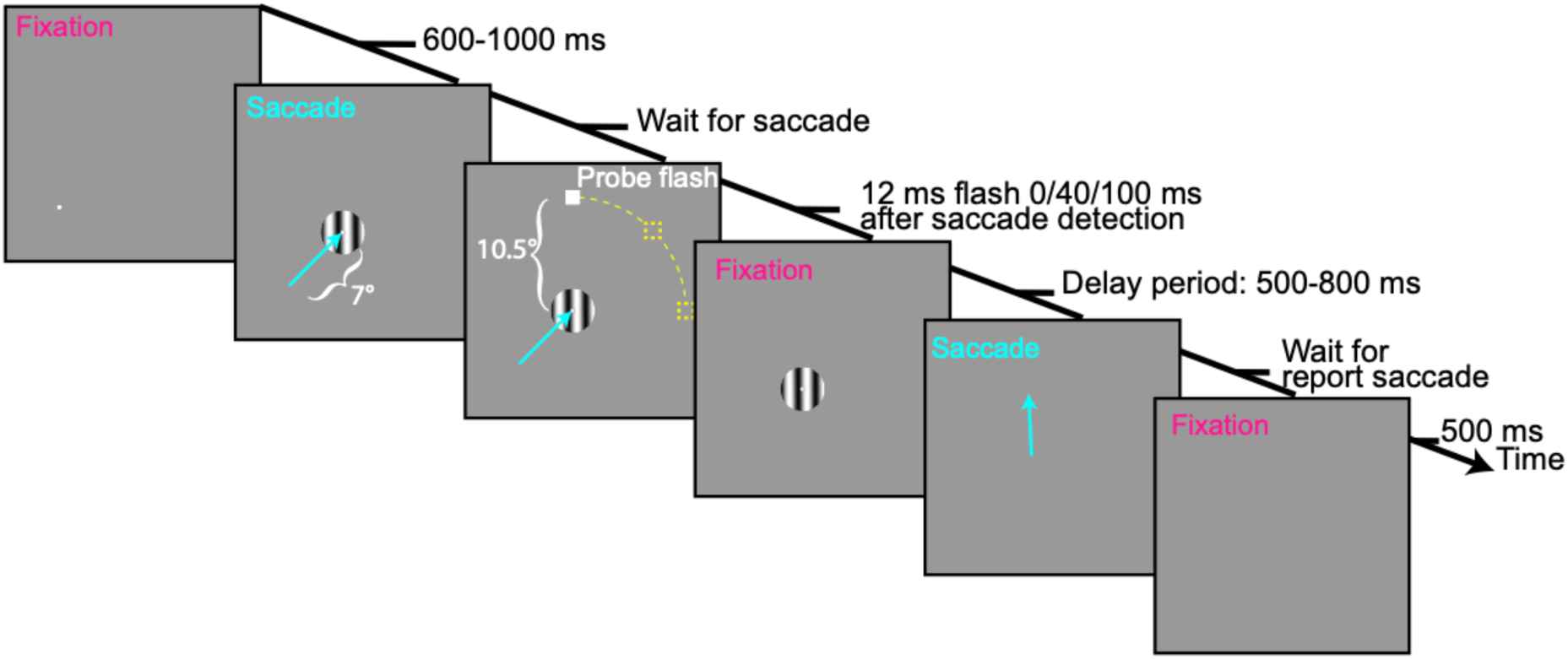
Perisaccadic visual mislocalization task for macaque monkeys. Each trial started with fixation of an eccentric spot (bottom-left corner of the display in the shown example). Upon removal of the spot, a saccade target appeared (Gabor grating), and the monkeys generated a visually-guided saccade towards it (oblique cyan arrow). We detected saccade onset online, and we presented a brief, single-frame probe flash at one of three possible delays from saccade detection. The probe flash could appear at one of three possible locations on a virtual circle (dashed yellow curve and placeholders) around the saccade target: one directly ahead of the saccade target along the saccade vector direction, and two at +/- 45 deg relative to the saccade vector (the shown example flash is at the 45 deg counterclockwise position). After an enforced delay, we removed the grating, and the monkeys reported their perceived flash location with a second, memory-guided, saccade. They had to maintain their gaze on the reported location for an additional delay time before being rewarded. In baseline trials, the probe flash appeared 100 ms after saccade detection and remained visible for 400 ms. Thus, it was clearly visible well after the end of the first saccade, resulting in an expected veridical percept.

We monitored the monkeys’ eye position in real-time and detected the onset of a saccade when the eye position deviated by 1.5 degrees from the initial fixation spot. Following saccade detection, a probe flash was presented either immediately or with a delay of 40 or 100 ms (in subsequent post-hoc analyses, we recalculated exact probe times; see below). The probe flash was displayed for a brief duration of ∼12 ms (1 frame at a display refresh rate of 85 Hz) and was positioned 10.5 degrees away from the center of the grating (7.5 deg for the slightly smaller cardinal saccades of monkey M). The probe flash could appear either directly along the saccade vector direction (and farther away from the saccade target) or at an angle of +/-45 deg relative to the saccade vector (again farther away than the saccade target location). Thus, there were three possible probe flash locations, all of which were expected to experience backward mislocalization for probe flash times very close to saccade onset, especially given the bright display and clear visual reference frames in the environment (Honda, 1999; Lappe et al., 2000).

After the monkeys made the visually-guided saccade, with a maximum allowed reaction time of 500 ms, they maintained stable fixation on the saccade target for an additional 500-800 ms; this delay period started once the eye entered a predefined window around the saccade target. After the delay period, the grating disappeared, cueing the monkeys to make a memory-guided saccade “reporting” the remembered probe flash location. The monkeys were rewarded if they made the report saccade within <500 ms, and we allowed a radius of 8 deg tolerance window around the true flash location for the landing position of the eye. This large tolerance window was intentional because we wanted to avoid penalizing the monkeys for their mislocalized percepts; otherwise, the monkeys would have learned, over time, to constrain their report saccades to the regions gaining them rewards. On successful trials, the monkeys received juice rewards; on unsuccessful trials (e.g. late initial saccade or broken fixation), the trials were aborted and juice rewards were withheld.

We also interleaved trials containing what we called a “baseline” condition. In this condition, which occurred in a total of 6.25% of all trials in a given session, the probe flash was always presented 100 ms after online saccade detection, and the probe also remained visible for a full 400 ms. That is, the probe was still clearly visible while the monkeys’ were in a stable fixation position on the grating center after the end of the primary saccade. This allowed assessing the monkeys’ eye movement behavior in the report saccade when no perceptual mislocalization was to be expected (of course, other small errors, such as those associated with memory-guided saccades, could still occur, as we also clarify in Results).

To avoid that the monkeys were solving the task by remembering absolute flash locations across trials, for example, by guessing their position relative to the edges of the display, we applied a global constant shift in all stimulus positions, which we changed randomly from trial to trial, but which we kept constant for all stimulus events within a trial. Specifically, on every trial, the whole stimulus geometry (initial fixation position, final fixation position, and the virtual circle on which the probe flashes could occur) was shifted randomly by a constant known amount, within a range of ±0.5 deg both horizontally and vertically. In all analyses, all positions were re-referenced to the true saccade target location on any given trial. This is a similar approach to our recent human perisaccadic mislocalization experiments (Baumann et al., 2024).

Across data collection sessions, we varied the primary saccade direction, in order to demonstrate the two-dimensional flexibility of our paradigm. We selected eight distinct initial fixation positions on a virtual circle around display center, allowing us to test eight different saccade vectors. Although our paradigm was designed to support any starting and end position in theory, we focused on these eight equally spaced directions for simplicity and for the opportunity to collect many trial repetitions of any given condition. In practice, we blocked trials of a given saccade direction in a given data collection session. This was done purely for pragmatic reasons. Specifically, we typically collected multiple sessions of data from each animal, and we wanted to have large numbers of trials for each saccade direction. Therefore, it was simpler to specify a given saccade direction for any given block that we ran. This is also the expected situation during neurophysiological experiments, in which a given saccade vector would be constant for a given recording site (for example, in the superior colliculus). However, once the monkeys are proficient in this task, we believe that randomizing saccade directions within a single session would still be very easily performed by them (see our note on training trajectory below).

For monkey M, as stated above, the cardinal saccades were only horizontal, and they had an amplitude of 5 deg (rather than 7 deg). Also, the probe flashes were 7.5 deg away from the saccade target in these cases (rather than 10.5). Thus, when summarizing this monkey’s results, we separated the oblique conditions (with 7 deg saccades) from the cardinal ones (see Results). We did this in order to avoid potential confusion with respect to the quantitative amplitude of mislocalization for the two different saccade sizes that we tested in this animal. On the other hand, showing qualitatively similar results with different saccade sizes, both in the same monkey and also across animals (see Results), further confirms our interpretation of the robustness and flexibility of our visual mislocalization paradigm.

Finally, in the task above, the report by the animals was made through a memory-guided saccade to the “perceived” flash location. Memory-guided saccades have systematic (and variable) errors in their landing positions (White et al., 1994; Willeke et al., 2019), which can also depend on saccade direction (Hafed and Chen, 2016). However, as mentioned above, and as we also show in Results, we varied saccade directions across our experimental sessions, in order to convince ourselves that what we saw was true mislocalization (Grujic et al., 2018) and not only errors in memory-based reporting (e.g. Fig. 4 in Results). Moreover, the perisaccadic flash eccentricities were far enough from the saccade target to make the peak mislocalization errors significantly larger than any systematic or variable errors expected from memory-guided saccades (see Results).

We collected approximately 350-500 successful trials per session from each monkey. In total, we analyzed 7582, 16005, and 20902 successful and accepted trials for the saccade direction analysis, as well as 14953, 23830, and 23371 successful and accepted trials for the saccade target appearance analysis from monkeys A, F, and M, respectively (see below for our definitions of saccade direction data analysis and saccade target appearance data analysis).

### Training trajectory

Success in our task was built on a very simple training trajectory that each monkey underwent, and that we describe here briefly. All three monkeys easily mastered the task.

All three monkeys were already experts in the individual task components before starting this project. Specifically, they knew how to perform both visually-guided and memory-guided saccades. To get the monkeys to report flash position in our current mislocalization task, we first started with the visually-guided saccade component of the paradigm; the monkeys simply made a saccade towards the grating image when they were instructed to do so (by removal of the initial fixation spot and the appearance of the grating). They were also rewarded classically, immediately after removal of the grating image.

We then introduced a second delayed, visually-guided saccade in every trial. Upon detection of the first saccade to the grating center, we presented the probe. However, instead of being flashed briefly, the probe remained visible until trial end. After the first saccade to the grating center, the monkeys performed a delayed saccade to the probe after grating removal. The reward now only came after a sequence of two instructed saccades (both visually-guided). Since the monkeys were already proficient in delayed, visually-guided saccades, this stage of training was achieved already from the first trial, allowing us to proceed.

We next introduced trials in which we presented the flash like in the main task above, except for two differences. First, the flash occurred always 100 ms after detecting the visually-guided saccade. That is, we were sure that it should be minimally mislocalized since it appeared much later than the saccade time (Ross et al., 1997; Grujic et al., 2018). Second, the flash was presented for a long period of time (400 ms) on every trial (like in our baseline condition). This ensured that the flash was minimally mislocalized by the previous saccade, and that it was still visible during stable fixation of the grating center. Upon removal of the grating, the monkeys made a relatively accurate memory-guided saccade towards the flash position, and we immediately rewarded them for having their gaze enter a relaxed virtual window of 8 deg radius around the true flash position. This made them quickly realize that the reward was now contingent on reporting the remembered flash position after completing their initial visually-guided saccade.

We then gradually increased the period in which the monkeys had to maintain fixation on the remembered flash position before getting rewarded. This allowed them to maintain stable gaze position for a few hundred milliseconds before getting rewarded (Fig. 1). Thus, by now, the monkeys were proficient in performing an initial visually-guided saccade to an image, and then making a second memory-guided saccade towards the remembered flash location (with the flash itself being veridically perceived since it occurred long after saccade onset, and also since it was visible for a long time). For one monkey (A), the animal had some difficulty transitioning to the memory-guided component of the task so far. Therefore, we introduced a few extra sessions in which the probe reappeared when the grating disappeared. This prompted the monkey to realize that the task was to focus on the remembered probe flash location.

When we were confident that the monkeys were performing this task with high fidelity, we then reduced the duration of the flash (down to a single display frame). And, finally, we started introducing the other flash times, in which we expected a large mislocalization error. As mentioned above, we maintained a relaxed virtual tolerance window radius around flash position, so that we did not penalize the monkeys for their mislocalized reports. All animals readily learned the task without much difficulties.

### Data analysis

#### Primary saccade processing

We detected all saccades (in post-hoc analyses) using the methods that we described previously (Chen and Hafed, 2013). For the primary saccades, such detection allowed us to recalculate probe flash times relative to the actual saccade onset in the data. We found that the probe flashes in the experiment occurred at three distinct times from saccade onset, consistent with our experimental design. Based on the recalculations, we specifically classified the probe flash times into the following three categories: probes occurring ∼30 ms after primary saccade onset, probes occurring ∼70 ms after primary saccade onset, and probes occurring ∼130 ms after primary saccade onset. That is, the online saccade detection process described above took approximately 30 ms to trigger the flashes during the data collection (expected given the dependence of our detection algorithm on eye position deviating sufficiently from initial fixation, and also given the frame refresh rate of the display). Figure 2A shows an example of the relationship between primary saccade times and probe flash onset times from one example session of monkey A. The figure plots radial eye velocity of the primary saccade (each curve is a saccade), with all trials aligned to probe flash onset time. There were three distinct time periods separating saccade and flash times across all trials.

**Figure 2.**
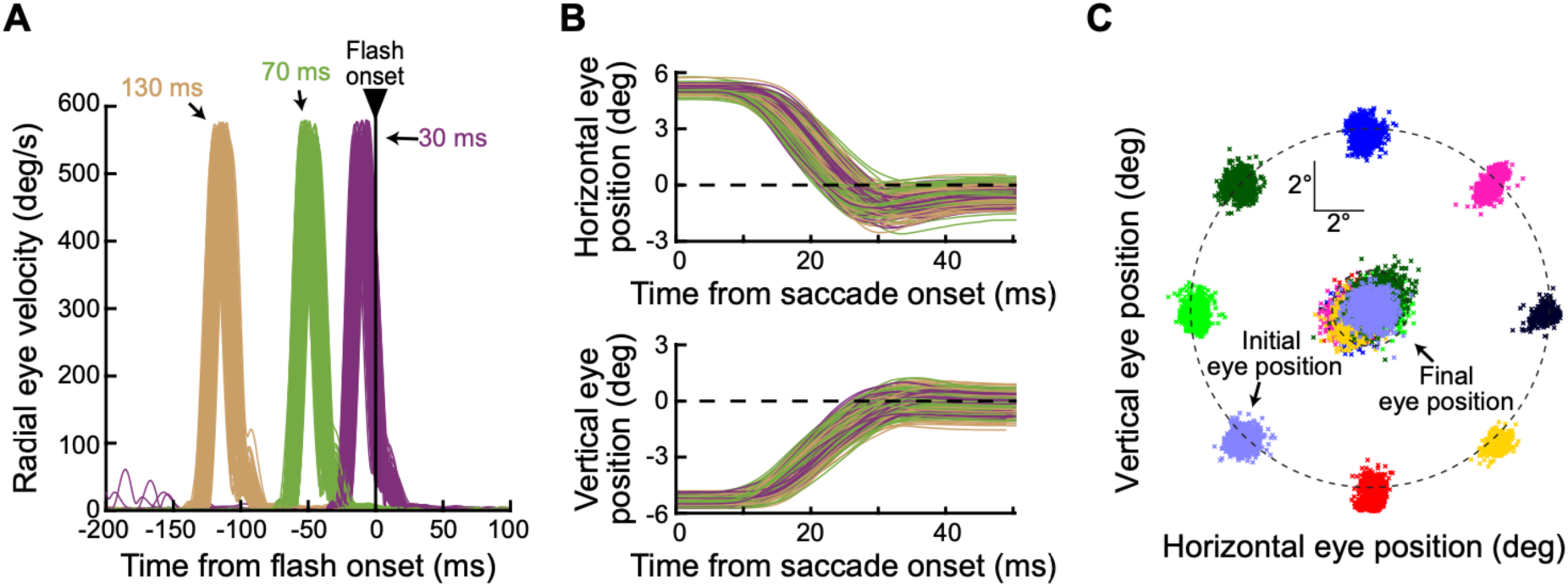
Controlling probe flash timing and saccade metrics and kinematics across conditions. **(A)** Radial eye velocity plots of all accepted primary saccades from an example session of monkey A. All traces are aligned to the onset of the probe flash, revealing three distinct perisaccadic flash times, which we categorized by the three different colors. We expected progressively less visual mislocalization for longer times between the saccades and the probe flashes. Baseline trials are not included in this panel. **(B)** Horizontal (top) and vertical (bottom) eye position traces from one example saccade vector direction (up and to the left) out of the same example session shown in A. Different colors show the different probe flash times, and the data suggest no systematic differences in the saccades across the flash times. **(C)** We also ensured that landing eye positions (central clouds of dots) overlapped for different saccade directions, again allowing us to investigate the influence of saccade direction on mislocalization strength with overlapping saccade properties (the kinematics were also matched as per the description in the text). The small black dashed circle indicates the extent of the grating towards which the primary instructed saccade was generated, and the big black dashed circle indicates the virtual circle along which the initial fixation positions were placed.

Besides checking timing, we also checked primary saccade consistency across conditions. That is, we wanted to avoid the possibility that any differences in visual mislocalization that we uncovered (for example, as a function of saccade direction) could be due to systematic differences in the actual primary saccade metrics and kinematics. Therefore, before analyzing the behavioral reports of the animals, we first filtered trials according to the eye movement data of the primary saccades. In one set of comparisons (see Results), we were interested in investigating the influences of primary saccade direction on perisaccadic visual mislocalization strength. Therefore, we ensured matched saccade vector and kinematic properties across the different directions of the primary saccades of our main task (Fig. 1). In another set of comparisons, we studied the influences of saccade target appearance on mislocalization strength; here, we matched the primary saccades across the different saccade target image types (low versus high spatial frequency).

For the vector and kinematic matching in the investigations of primary saccade direction influences, we binned all saccade main sequence data (Zuber et al., 1965; Bahill et al., 1975) into intervals of 0.5 deg in amplitude by 40 deg/s in peak velocity. We included in the analyses only those trials from the main sequence bins that had at least one trial from each of the eight saccade directions (four in the case of monkey M’s oblique saccade data; this monkey’s cardinal saccades were analyzed separately due to their smaller size). This ensured that we could compare saccades across different directions having similar ranges of amplitudes and peak velocities.

We also noticed that monkey M had a substantial difference in saccade peak velocity between rightward and leftward saccades (∼350 versus ∼550 deg/s, respectively). Therefore, for this monkey, we decided to report all analyses separately according to the hemifield of the saccade vector (see Results), meaning that we also separated the vector and kinematic matching procedures in this monkey for each hemifield individually. This way, we did not mask any possible quantitative effects of saccade peak velocity on mislocalization amplitude, but the results were always qualitatively the same across hemifields (and across animals).

Figure 2B shows, for the same monkey and session as in Fig. 2A, example accepted saccades. Each panel shows either horizontal (top) or vertical (bottom) eye position aligned on saccade onset, and the different colors show the different probe flash time classifications of Fig. 2A. Figure 2C, in turn, shows all start and end eye positions from all trials of all sessions of the same monkey. These trials passed the vector and kinematic matching (for the investigation of saccade direction influences on visual mislocalization), and they, therefore, went into further analysis.

For investigating the influences of saccade target appearance on mislocalization strength, we applied the same binning strategy above in order to achieve vector and kinematic matching, but we now compared saccades to either low or high spatial frequency Gabor gratings. We, thus, pooled across all saccade directions. We included only those trials from the main sequence bins that contained at least one trial repetition from each of the two saccade-target image conditions (low and high spatial frequency) of each probe flash time. This ensured that saccades were matched for both metrics and kinematics across the different image conditions of the saccade target.

#### Establishing the existence of perisaccadic mislocalization

We next analyzed the report saccade endpoints, which constituted our experimental measure of perisaccadic visual mislocalization. We had three general classes of analyses in the study. First, we established that we were indeed measuring genuine mislocalization and not artifacts of either memory-guided saccade errors or our particular analysis choices. Second, we focused on the effects of saccade direction on mislocalization properties. Finally, we investigated the effects of saccade target appearance. We now describe the first class of analyses.

To establish that we were measuring genuine mislocalization, we took the following measures, including control analyses to rule other potential explanations out. For each flash time and saccade direction, we measured the final eye position of the report saccade on every accepted trial, and we then plotted the raw two-dimensional data (e.g. Fig. 3D in Results) relative to the geometry of the saccade vector and probe flash location. We also collected summary statistics within each monkey. Specifically, for each probe flash location and time (for a given saccade direction), we calculated the mean and SEM of all reported probe flash locations. This allowed us to visualize the pattern of mislocalization (e.g. Fig. 3E, left in Results).

**Figure 3.**
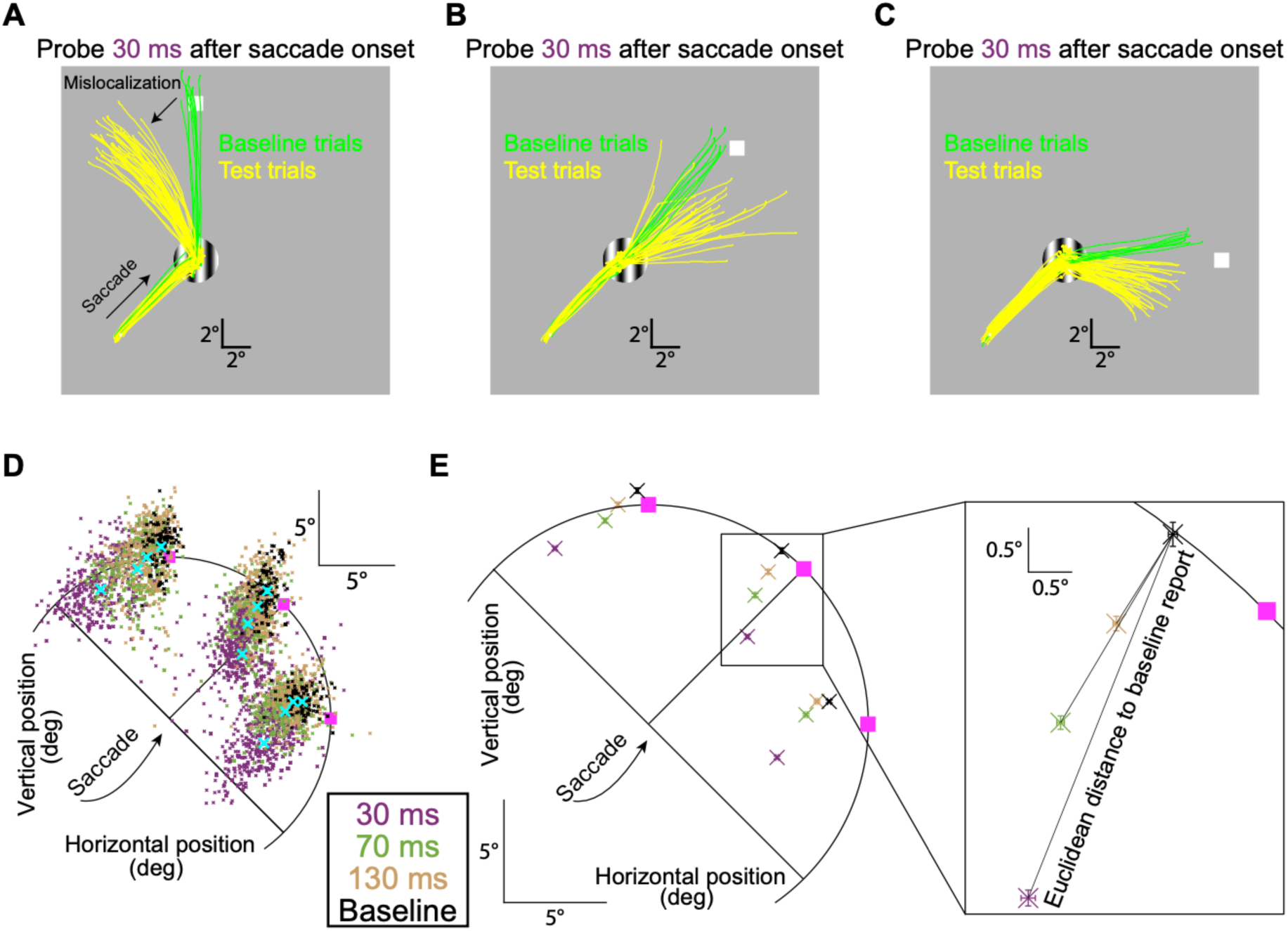
Two-dimensional perisaccadic “compression” in rhesus macaque monkeys. **(A)** Example eye position traces from monkey F when generating an oblique (upward/rightward) primary saccade, followed by a second report saccade. Individual traces show the entire eye movement sequence per trial (Materials and Methods), and we selected only trials with probe flashes directly above the saccade target location. The probe flash is schematized with a white square, but it was not actually visible at the time of the second saccade (the grating could also be different across trials; Materials and Methods). Green traces show baseline trials in which the probe flash remained visible for a prolonged period after the end of the primary saccade, but it was still invisible at the report saccade time (Materials and Methods). The monkey’s report saccade was largely accurate, with a small systematic error expected from memory-guided eye movements. For perisaccadic flashes (yellow traces), there was a strong distortion in the opposite direction from the saccade vector (compare black oblique arrows). **(B, C)** Example trials from the same monkey for two other flash locations. There was always reverse mislocalization. **(D)** Across all sessions from the same monkey, and for all probe flash times and the same oblique saccade direction, we plotted the two-dimensional reported locations. On baseline trials (black), the percepts were mostly veridical (with expected small, but systematic, errors associated with memory-guided saccades). For perisaccadic flashes (colored small dots), there was maximal mislocalization at 30 ms, with a gradual temporal recovery. This is consistent with human mislocalization time courses. The pink squares indicate true flash locations, and the cyan crosses show the means of each colored cluster (also presented with proper color coding in **E**). **(E)** The main plot (left) shows mean and SEM summaries of the data of **D**. For longer and longer times, gradual recovery towards baseline performance occurred at each probe flash location. On the right, we magnified the data for one probe flash location, to demonstrate our summary measures in subsequent figures: we characterized mislocalization strength by calculating the Euclidean distance between reports on mislocalization trials to reports on baseline trials (Materials and Methods). In separate control analyses, we also checked the errors in baseline reports themselves (from true flash location).

We quantified mislocalization strength as follows. For each flash location, we used the baseline condition to know the average reported location by the monkey when perception was expected to be veridical. Then, for each trial of the same flash location but for one of the experimental perisaccadic flash times, we measured the Euclidean distance between the reported location of this trial and the mean reported location of the baseline condition (e.g. Fig. 3E, right in Results). This allowed us to obtain average mislocalization across trials of a given flash time relative to any systematic error that might have existed in the report saccade in the absence of any perisaccadic mislocalization.

To explicitly confirm that we did indeed need to account for systematic errors in memory-guided saccades (White et al., 1994; Willeke et al., 2019), we also directly compared the error patterns between baseline and mislocalization trials relative to true flash location. For each trial (whether baseline or with a brief perisaccadic flash), we computed an error vector by connecting the veridical flash location to the reported location. We then calculated the angle of this vector relative to the primary saccade direction angle. If mislocalization errors were systematically related to the primary saccade direction, then their angular distributions were expected to be very tightly distributed around 180 deg (i.e. backward mislocalization). However, on the baseline trials, if the errors were just systematic errors devoid of any perisaccadic mislocalization, then they were not expected to depend on the primary saccade direction; thus, the angle distributions relative to the primary saccade direction were expected to span 360 deg.

We also confirmed that baseline reports were always more veridical than the reports for perisaccadic flashes, and we verified that differences in baseline errors (e.g. as a function of saccade direction) were not sufficient to explain the differences in peak mislocalization effects that we reported in Results.

Having established that the baseline trials were the proper reference measure for estimating veridical percepts, we next considered whether memory decay could account for the larger localization errors observed in perisaccadic trials than in baseline trials (which we report in Results). Specifically, the longer flash durations in our baseline trials (relative to perisaccadic test trials) resulted in shorter delays between the probe flash offset time and the go-signal for the animal to report the probe flash location. To confirm that this was not sufficient to explain the differences in error magnitudes between perisaccadic and baseline trials, we explicitly related localization error amplitude in baseline trials to delay period duration. We binned delay period durations into 30 ms bins, and we plotted the baseline report error amplitudes as a function of these bins. This allowed us to assess whether error amplitudes on perisaccadic probe flash trials were simple extrapolations of memory decay effects.

We then examined the relationship between mislocalization strength and flash onset time relative to saccade offset, by implementing a more fine-grained temporal binning approach. Within each of the three original coarse flash time conditions, we introduced additional temporal bins to increase resolution. This finer binning was made possible by the natural variability in saccade durations across trials, and it allowed us to confirm that we were likely not observing a biphasic mislocalization pattern as might be expected from pure darkness conditions (Honda, 1989, 1991, 1993, 1999).

To ensure that the observed mislocalization effects were also not merely a consequence of differences in eye position during (intrasaccadic and postsaccadic) probe flash presentations, we assessed the relationship between eye position at the time of the probe flash onset and the mislocalization strength. For each monkey and each probe flash time condition, we binned the mislocalization strength according to the distance between the eye position and the saccade target at the moment of the probe flash presentation. Bins were set at intervals of 0.2 degrees. Eye positions were assigned negative values when the eye was located between the initial fixation point and the saccade target, and positive values indicated overshoots beyond the saccade target. This analysis allowed us to confirm that mislocalization magnitudes depended on flash times from primary saccade onset rather than on the exact eye position at probe flash occurrence (see Results).

On a related note, we next validated our expectation that differences in mislocalization strengths across probe flash presentation times were not driven by the presence of corrective saccades during the delay periods. We analyzed the temporal distribution of corrective saccades relative to the initial saccade onset. We aligned corrective saccade onset time to the initial saccade onset time, in order to assess these saccades’ timing across different probe flash time conditions.

#### Exploring the effects of saccade direction on perisaccadic mislocalization

After being convinced that we were measuring genuine perisaccadic mislocalization, we proceeded to summarizing the effects of saccade direction on visual mislocalization strength. In one analysis, we averaged the Euclidean distance measure described above across all trials that involved either upward or downward saccades for each probe flash time, respectively. Upward saccades in this kind of analysis included oblique upward/rightward and oblique upward/leftward saccades, in addition to purely vertical upward saccades. Similarly, downward saccades also included the oblique saccades with a downward component. In this comparison, we averaged across all three probe flash locations of a given saccade direction. In related analyses, we also measured the mislocalization strength (Euclidean distance to the reports in the baseline condition) for each saccade direction separately. Again, we pooled all flash locations per saccade direction together in this analysis, but we always treated flash times separately. Of course, we had previously inspected all the raw data of individual flash locations (e.g. Figs. 3D, 5A, B in Results) to justify pooling them in our summary analyses.

We also further summarized the difference in mislocalization strength between upward and downward saccades in a given analysis. To do so, we calculated the mathematical difference (at any given flash time) between the Euclidean distance measure (to the respective baseline condition of each flash location) obtained for upward saccades and the Euclidean distance measure obtained for downward saccades. We then plotted this difference measure across flash times. Again, this is a similar approach to how we reported changes in mislocalization strength across different experimental conditions in our recent human work (Baumann et al., 2024).

To once more validate that our observations were not dependent on our use of the baseline reports, we also repeated our saccade direction comparisons but by now relating the mislocalization errors to the reports on the trials with flashes presented ∼130 ms after saccade onset (rather than the baseline trials). That is, we exploited the fact that perception recovered with long times after saccade onset, and used the latest flash time as the condition with closest-to-veridical percepts. As expected, similar results were observed.

Since perisaccadic mislocalization for a given saccade direction can also have a two-dimensional landscape (Kaiser and Lappe, 2004; Grujic et al., 2018), we additionally investigated the vector components of mislocalization, similar to what we had done recently with humans (Baumann et al., 2024). We examined the component of localization error that was parallel to the saccade direction and the component that was orthogonal. This analysis followed the same approach as that described above (measuring Euclidean distances to baseline reports for the different saccade directions), except that we separated the parallel and orthogonal components of the mislocalization errors.

In a separate comparison, we focused solely on the trials with horizontal primary saccades. In this case, we compared the Euclidean distance measure for probe flash locations in the upper versus lower visual fields. Thus, for such horizontal saccades, we compared the mislocalization strength between the two oblique probe flash locations associated with these saccades (by definition of the task, one oblique probe flash location would have been in the upper visual field, and ahead of the saccade target location, and the other would have been symmetrically in the lower visual field). We should also note here that even for vertical saccades, probe flashes ahead of the saccade target could be in either the upper or lower visual field (e.g. upper visual field for upward saccades and lower visual field for downward saccades). For oblique saccades, some flash locations were at the same vertical eccentricity as the saccade target, but they were still in the upper visual field presaccadically for upward oblique saccades, and still in the lower visual field presaccadically when the oblique saccade was downward. Thus, it is likely that some of our saccade direction effects interacted with visual field effects, as was also previously observed in humans (Grujic et al., 2018).

#### Exploring the effects of saccade target appearance on perisaccadic mislocalization

We also performed an investigation of the influences of the saccade target appearance on perisaccadic visual mislocalization. To do so, we averaged the Euclidean distance measures for all saccade directions and probe flash locations, but now separated trials based on the spatial frequency of the saccade target.

#### Statistical analyses

We statistically compared different conditions using a two-way ANOVA on the Euclidean distance measure, with the two factors of the ANOVA being probe flash time relative to saccade onset and either saccade direction, flash location, or saccade target appearance, depending on the particular comparison that we were making. We always performed these analyses within each monkey individually, to be able to compare the outcomes across the individual animals. Although we consistently presented monkey M’s leftward and rightward saccades separately in all figures, we did not distinguish between them in the statistical analyses, as this separation did not affect the statistical results. We also statistically compared baseline reports under different comparison conditions. For example, we checked whether the errors in baseline reports (relative to the true flash location) were different whether saccades were upward or downward, and we did so by performing (for each monkey) a t-test. Details are provided in the relevant sections of Results, and the statistics were performed with IBM SPSS (Version 29.0).

## Results

We aimed to establish a robust and flexible visual mislocalization paradigm in rhesus macaque monkeys. We tested this paradigm on three animals, and they all exhibited clear perisaccadic mislocalization, with expected recovery time courses. These animals also all replicated prior human findings of stronger mislocalization with upward saccades (Grujic et al., 2018), as well as stronger mislocalization for upper visual field flashes (with horizontal saccades) (Baumann et al., 2024). We also found subtle effects of saccade target image appearance on mislocalization strength (Baumann et al., 2024), but these effects were not as consistent as with humans, and they were also not as strong as the effects of saccade directions and flash locations in the same animals. We summarize all of these observations below.

### Monkeys reported perisaccadic flashes as being closer to the saccade target

We trained our monkeys to report the location of a brief (∼12 ms) probe flash presented at different perisaccadic time points (Figs. 1, 2). They did so by directing their gaze, via a memory-guided saccade, to the remembered flash location after an instruction to do so (Fig. 1). Note that such memory-guided reporting of percepts is present even in classic human invocations of this same task (Ross et al., 1997; Lappe et al., 2000). To confirm that there was indeed mislocalization, we interleaved baseline trials in which the flash was long-lasting, and thus clearly visible after primary saccade end (Materials and Methods). Therefore, any errors in reporting for this baseline flash condition should have been restricted to only those errors (whether systematic or variable) associated with memory-based behavioral reports (White et al., 1994; Willeke et al., 2019; Willeke et al., 2022), and not directly related to perisaccadic mislocalization.

As a first step in our analyses, we first performed a detailed investigation of unsuccessful trials that were not analyzed for mislocalization reports. We did this in order to gain further insights into the monkeys’ task performance and potential strategies. Unsuccessful trials were defined as those in which the monkey correctly initiated the trial by performing the instructed primary saccade (thereby triggering the probe flash presentation) but subsequently failed to obtain the reward due to performance-related errors. Trials in which the monkey did not initiate the task at all (e.g., due to a lack of motivation, blinks, or eye position instabilities at the initial fixation spot) were excluded from this analysis. Across all conditions, all three monkeys successfully completed 84.4%, 79.3% and 76.7% (Monkeys A, F, and M, respectively) of the initiated trials, confirming that the probe flash was clearly visible under all experimental conditions. The failure analysis revealed two primary categories of unsuccessful trials: (1) fixation breaks, and (2) incorrect saccade landings. Fixation breaks were errors in which the monkey broke fixation either before the go signal for the report saccade or during the final fixation period after the end of the report saccade. The proportion of fixation breaks was 9% for Monkey A, 12.5% for Monkey F, and 10.7% for Monkey M. Incorrect saccade landings were errors in which the monkey executed the report saccade but failed to land within the predefined spatial window required for successful reporting. The rates of this error type were 6.6% (shortest flash delay: 4.9%, medium delay: 1%, longest delay: 0.5%, and baseline: 0.2%) for Monkey A, 8.2% (shortest delay: 5.7%, medium delay: 1.3%, longest delay: 0.8%, and baseline: 0.4%) for Monkey F, and 12.5% (shortest delay: 9%, medium delay: 2%, longest delay: 0.5%, and baseline: 1%) for Monkey M. Thus, landing errors were always more frequent for the shortest probe flash times relative to the primary saccade onset.

The increased error rates for short probe flash delays are already the first hint that the monkeys really did experience perisaccadic visual mislocalization. Specifically, based on human research, shorter delays between saccade onset and probe flash presentation are associated with the largest mislocalization effects. Given the fixed spatial tolerance windows that we employed for accepting correct responses, larger mislocalization errors increased the likelihood that the monkeys’ reported locations fell outside of these windows, leading to a higher task failure rate. In essence, when the perceived location of the probe was strongly shifted, the monkeys might have been aiming accurately based on their perception, but the degree of mislocalization caused their responses to be registered as failures. This observation encouraged us to further analyze the successful trials for hallmarks of perisaccadic mislocalization. All results that follow are now solely based on the successful and accepted trials.

Figure 3A shows example results from monkey F when generating an oblique upward/rightward primary saccade. The probe flash location in this case was rotated 45 deg counterclockwise from the saccade vector direction. In the figure, the probe flash is indicated by a white square, but it was not actually visible at the time of the report saccade. Each trace in the figure shows eye position from one example trial, and different traces show different trials. We plotted example trials from the baseline condition in green, and example trials from a perisaccadic condition (the one with the probe flash occurring 30 ms after saccade onset) in yellow. As can be seen, the primary saccades (oblique upward/rightward movements and directed towards the grating center) were all the same whether the trials were baseline trials or perisaccadic flash trials; that is, the eye position traces nicely overlapped with each other. This is expected given our eye movement controls mentioned earlier (Materials and Methods). Importantly, the report saccades were very different depending on the properties of perisaccadic visual stimulation. In the baseline trials, the report saccades had a veridical direction (upward in this case); for brief perisaccadic flashes occurring 30 ms after saccade onset, the reported locations were strongly shifted backward along the reverse direction of the primary saccade vector (compare the black oblique arrows). Note that the baseline saccades were still memory-guided, just like the report saccades in the perisaccadic test trials. Therefore, the large difference between report saccades in the two conditions in Fig. 3A was attributed to brief, perisaccadic visual stimulation in one of the conditions, and not due to the memory-guided nature of the report saccades. Also note that there was larger variance in the report saccades in the perisaccadic test condition; this is also true in human perisaccadic mislocalization studies (Baumann et al., 2024). Thus, this monkey experienced strong backward perisaccadic mislocalization for brief flashes presented 30 ms after saccade onset, and this mislocalization happened with an oblique (rather than cardinal) saccade.

We also checked the other probe flash locations associated with this same saccade direction. Example traces from the same monkey and session are shown in Fig. 3B, C. Once again, there was always reverse mislocalization opposite in direction from the saccade vector. Specifically, in Fig. 3B, the probe flash was directly ahead of the saccade target, and the reported locations were generally in the same direction as the flash but closer to the saccade target location (mislocalized backward in direction). Similarly, with the probe flash location rotated clockwise relative to the saccade vector (Fig. 3C), the reported locations were now deviated both leftward and downward relative to the baseline reports (resulting in a mislocalization vector directly opposite the upward and rightward vector of the primary saccade). Thus, the mislocalization that this monkey experienced perisaccadically was always backward relative to the saccade vector direction, regardless of flash location.

Note also that there were systematic errors in the baseline report saccades themselves in Fig. 3A-C, which are expected (White et al., 1994; Willeke et al., 2019). For example, in Fig. 3C, a pure rightward report saccade was associated with a slight upward bias in eye position endpoints, along with saccade hypometry (White et al., 1994; Willeke et al., 2019; Willeke et al., 2022). However, the perisaccadic mislocalization errors were always larger than these baseline report errors. Moreover, the perisaccadic mislocalization errors had a clear systematic relationship in direction relative to the saccade vector direction, as we also describe in more detail below. Thus, the perisaccadic mislocalization effects of Fig. 3A-C were not fully explained by systematic errors associated with the reporting process itself (also see Fig. 4 below for a more detailed analysis of baseline reports).

**Figure 4.**
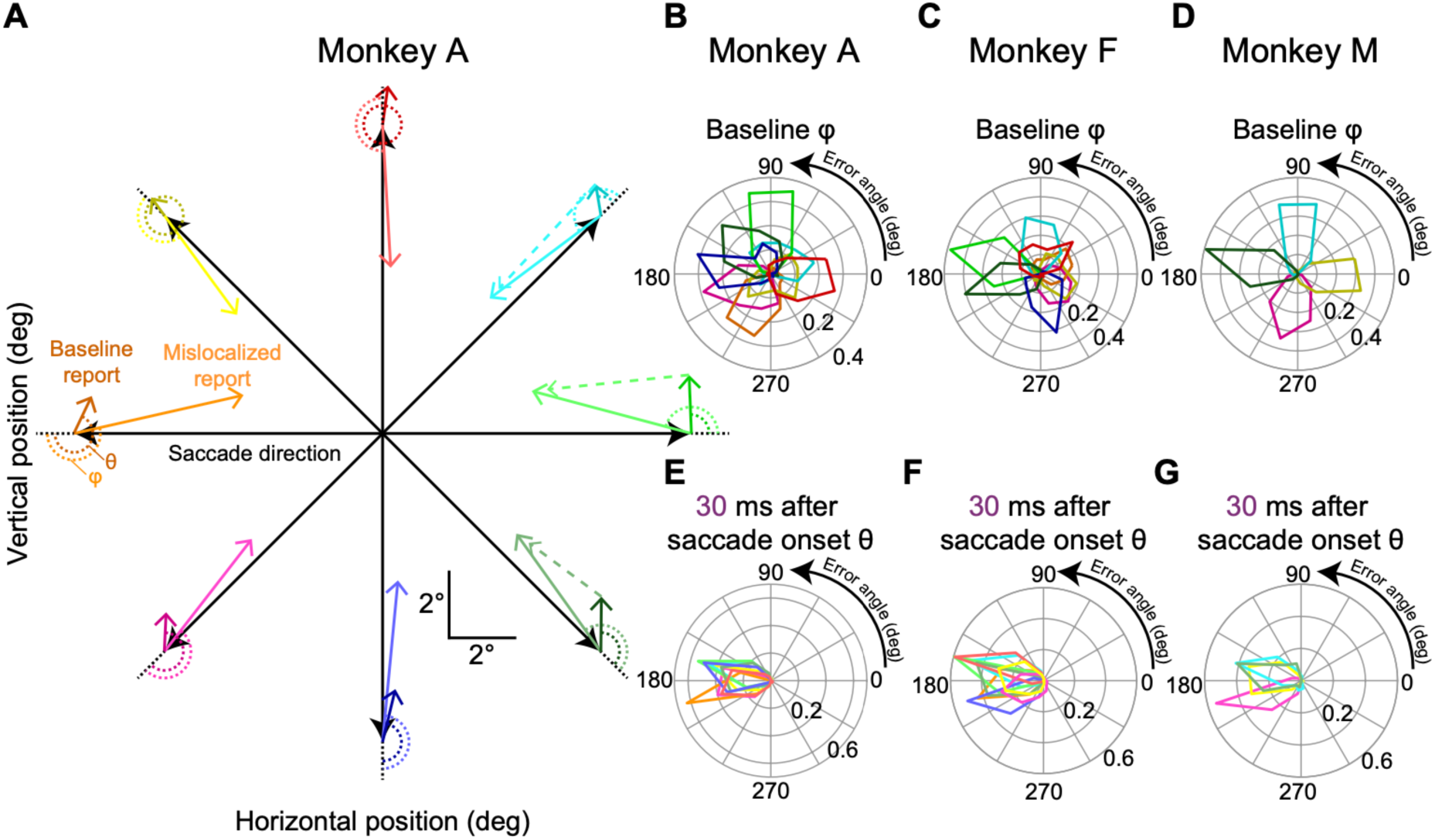
Comparison of error vector patterns between baseline and perisaccadic mislocalization trials. **(A)** Visualization of error vectors for Monkey A during the first flash time condition (30 ms after saccade onset), when mislocalization errors were most pronounced. Each colored vector connects the veridical flash location to the reported location, illustrating both baseline (darker-colored arrows) and mislocalization (brighter-colored, solid arrows) error vectors relative to the primary saccade direction (black arrows). Baseline errors consistently showed a small upward bias, independent of saccade direction. In contrast, mislocalization errors were systematically oriented in the direction opposite to the saccade vector. Dashed arrows indicate the direction of the mislocalization vector relative to the baseline report endpoint for selected saccade directions, demonstrating an even stronger 180-degree relationship with the primary saccade direction (other dashed vectors were not shown just to avoid clutter). **(B-D)** Polar histograms of angular errors (φ, as defined in **A**) for baseline error vectors across all primary saccade directions for Monkeys A, F, and M. Baseline errors showed a broad distribution spanning all 360 degrees, highlighting their independence from the saccade direction. Moreover, for each saccade direction, the baseline errors were clustered together, suggesting a single systematic baseline error direction independent of primary saccade direction (as in **A**). **(E-G)** Polar histograms of angular errors (θ, as defined in **A**) for mislocalization error vectors for Monkeys A, F, and M. Mislocalization errors were tightly clustered around 180 degrees relative to the saccade direction, indicating a robust and systematic divergence from baseline error patterns. **Extended Data Figure 4-1** shows the relationship between baseline localization errors and the delay period preceding report saccade execution. **Extended Data Figure 4-2** illustrates the relationship between mislocalization strengths and flash onset times relative to saccade offset, providing a more detailed time course depiction of perisaccadic mislocalization. **Extended Data Figure 4-3** demonstrates a lack of influence of eye position at the time of the probe flash presentation on mislocalization strength. **Extended Data Figure 4-4** shows the timing of corrective saccades relative to initial saccade onset across different probe flash presentation times.

Across all trials and all probe flash times and locations for the same oblique saccade direction, we found systematic mislocalization that gradually recovered with time from saccade onset. This is shown in Fig. 3D. Different colors indicate the different probe flash times, and individual small data points are individual trials (the cyan crosses show the means of each cluster). There was gradual recovery in mislocalization towards the baseline report locations when the probe flash time increased. This systematic trend can also be seen in Fig. 3E (left), in which we plotted the mean and SEM summary statistics of the same data. In the same figure (Fig. 3E, right), we also explained our measure of mislocalization strength that we used in subsequent figures. Specifically, mislocalization strength was defined as the Euclidean distance to the baseline report location for each probe flash position (Materials and Methods).

That the baseline reports were a proper reference point for canceling systematic errors of memory-guided saccades became even more clear to us when we related these baseline reports to individual saccade directions. Specifically, we compared error patterns in baseline trials to those observed during perisaccadic mislocalization (for the first flash time in which mislocalization was expected to be strongest from Fig. 3). The results of this analysis are presented in Fig. 4. Figure 4A shows (for monkey A) each primary saccade direction as a black arrow. For each such direction, we also represented the vector of the average baseline report error as a dark-colored arrow, and the vector of mislocalization (for the shortest flash time) as a ligher-colored arrow of the same general hue. As can be seen, mislocalization vectors were always opposite the saccade vector, but baseline error vectors were always of the same size and direction independent of primary saccade direction. Consistent with this, measuring the angle of the baseline error vector relative to primary saccade direction resulted in distributions spanning 360 deg (Fig. 4B-D, with each panel showing each monkey). On the other hand, measuring the angle of the mislocalization report vectors relative to primary saccade direction always revealed a 180 deg relationship (Fig. 4E-G). Thus, our mislocalization reports (Figs. 3, 4) were very distinct from baseline errors. Most importantly, closer inspection of Fig. 4A revealed that the baseline errors were (expectedly) always present, even in the mislocalization reports themselves. For example, for three sample primary saccade directions (those with a rightward component in the figure), when we corrected for the systematic baseline report vector error, mislocalization reports were better aligned with the 180 deg axis relative to the primary saccade direction than before. This makes perfect sense: if memory-guided saccades have systematic errors, then these systematic errors will still be present on brief probe flash trials. Thus, it was a valid choice for us to use baseline reports as the reference for veridical percepts in our analyses.

Our results so far suggest that we replicated human evidence of reverse perisaccadic visual mislocalization. Importantly, we did so for a non-horizontal saccade direction (Grujic et al., 2018), paving the way for generalized use of this paradigm in neurophysiological experiments. However, before diving deeper into additional properties of perisaccadic visual mislocalization in macaque monkeys, we first ruled out some additional caveats that might come up, especially with respect to our specific reporting mechanism (memory-guided saccades).

First, our baseline trials had a particularly short time between probe flash onset time and the go signal for reporting (due to the long duration of the probe flash on these trials; Materials and Methods). Thus, it might be wondered whether our larger mislocalization effects in Figs. 3, 4 (for the non-baseline trials) were due to decays in working memory associated with longer wait times on perisaccadic probe flash trials. To assess this, we analyzed the relationship between delay period duration after probe flash offset and localization accuracy on the baseline trials (Extended Data Figure 4-1). Even though there was a very subtle increase in localization error with longer delay durations, the effect was minimal and, even when extrapolated, it could not explain the mislocalization error amplitudes observed for the shortest flash onset times on the perisaccadic test trials. These findings align well with previous studies on memory-guided saccades, which demonstrated only minor (if any) memory decays over extended delays (White et al., 1994; Willeke et al., 2022). Thus, by including baseline trials in our analyses, we accounted for any general memory-related systematic errors, which were very temporally stable, ensuring that the reported mislocalization effects predominantly reflected genuine perisaccadic perceptual phenomena.

**Extended Data Figure 4-1.**
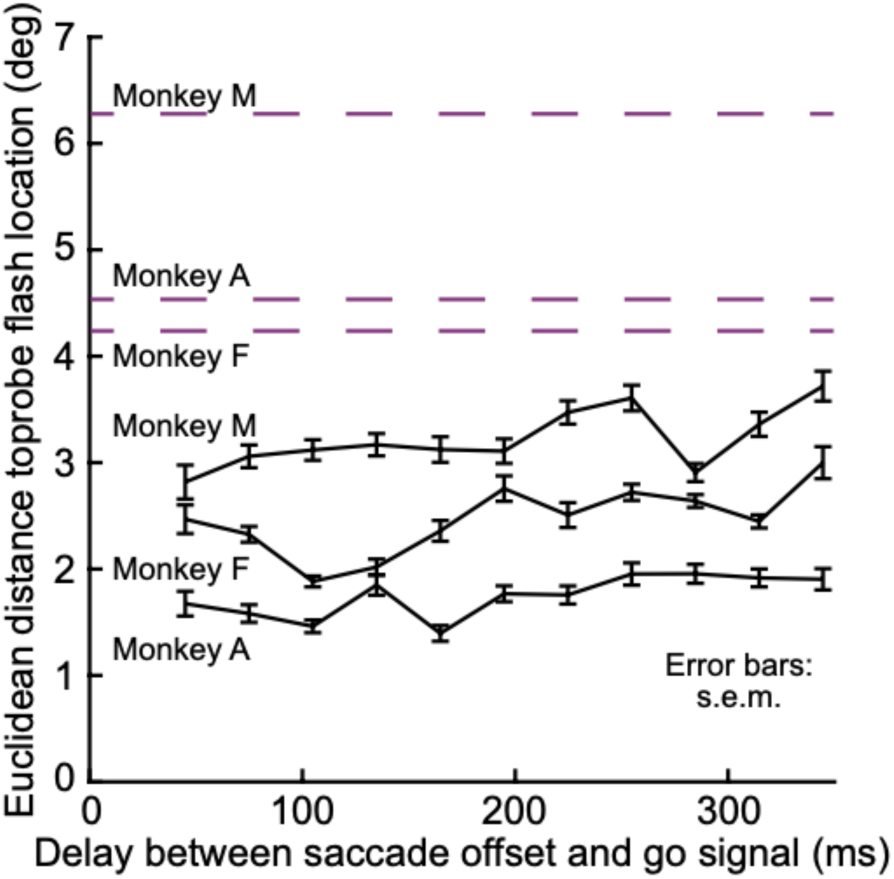
Relationship between delay period duration and baseline localization error. Euclidean distance between the reported location and the actual probe flash position in baseline trials as a function of delay duration between primary saccade offset and the go signal for the report saccade. Each data point represents mean localization error per time bin, with error bars indicating SEM. The dashed purple lines indicate the peak mislocalization values for the individual monkeys, providing a reference for the maximal observed perceptual distortions in the perisaccadic period. Even with extrapolation for much longer delays, the baseline report errors were much smaller than actual perisaccadic mislocalization strengths.

To better contextualize the temporal dynamics of mislocalization, we also examined the relationship between mislocalization strength and the onset time of the probe flashes relative to saccade offsets. Aligning the data on saccade offsets enabled more fine-grained temporal analyses due to saccade duration variability. Extended Data Figure 4-2 shows the time courses for individual monkeys, revealing a continuous decrease in mislocalization strength over time, indicative of a gradual recovery.

**Extended Data Figure 4-2.**
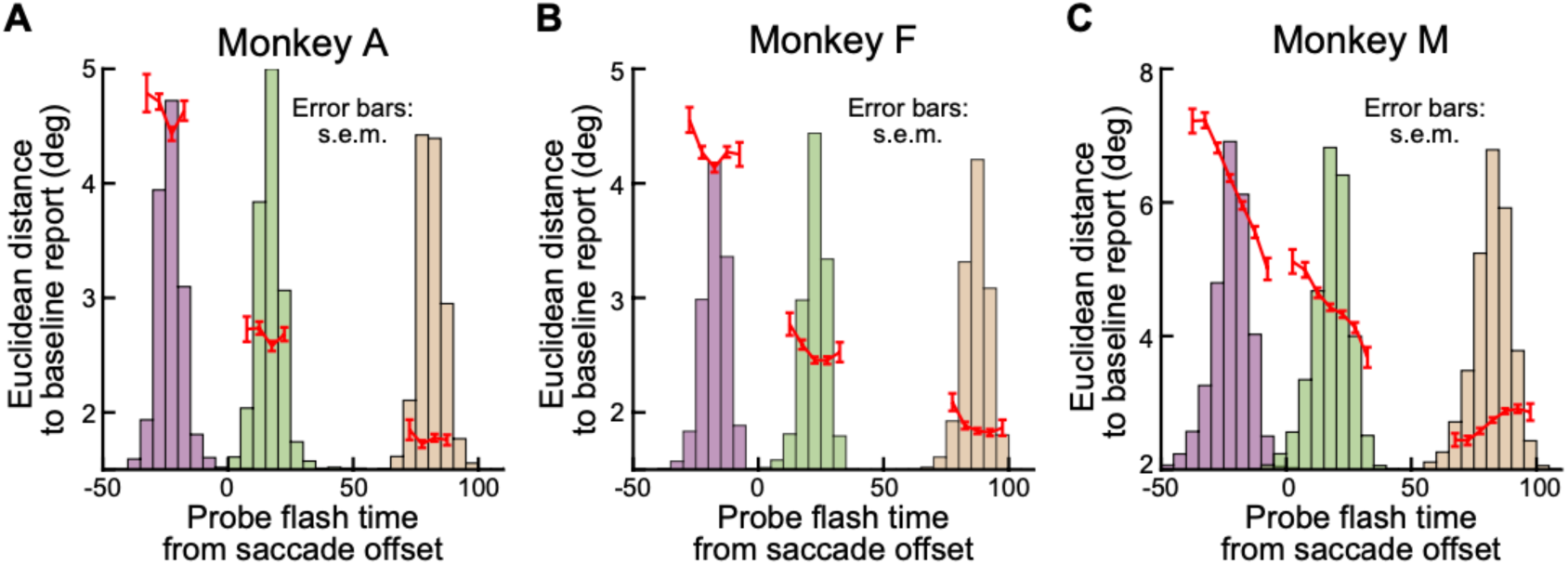
Time course of perisaccadic mislocalization relative to saccade offset. **(A-C)** Euclidean distance to the baseline report as a function of probe flash onset time relative to saccade offset for each monkey (red). Mislocalization gradually decreased with increasing delay after saccade offset, indicating a recovery of veridical localization over time. Error bars represent SEM. The faint histograms in the background show the actual underlying probe flash times; note that we did not plot red data points if there were fewer than 100 observations in any histogram bin.

Because eye position could be variable at probe flash time, especially for intrasaccadic probes (shortest flash time), we also ruled out eye position effects as an explanation for the strong mislocalization results that we saw so far (e.g. Figs. 3, 4, Extended Data Figure 4-2). We plotted the distance of the eye position from the saccade target at the moment of the probe flash presentation against mislocalization strength for each monkey and probe flash time (Extended Data Figure 4-3). Across all three animals, no systematic relationship was observed between eye position at flash time and mislocalization strength. For instance, although the distributions of eye positions for probe flash times of 70 and 130 ms after saccade onset were very similar (as the saccade had already concluded for both flash times), a substantial difference in mislocalization strength persisted between these two conditions. This suggests that probe flash timing relative to the primary saccade exerted a much stronger influence on localization performance than the precise eye position at the time of flash presentation.

Finally, corrective saccades following the primary saccade also did not account for the observed differences in mislocalization strengths over time. In Extended Data Figure 4-4, we analyzed the temporal distribution of corrective saccades relative to the primary saccade onset. As expected from saccadic refractory periods, nearly all corrective saccades occurred after the probe flash presentations. Moreover, the timing of corrective saccades was highly similar for flash times of 30 and 70 ms after saccade onset. For trials with probe flashes presented 130 ms after saccade onset, corrective saccades were slightly delayed. This delay is due to saccadic inhibition, where the probe flash transiently suppresses the execution of corrective saccades when presented in close temporal proximity (Buonocore and Hafed, 2023). The presence of saccadic inhibition further supports the idea that the monkeys attended to the probe flash (White and Rolfs, 2016).

**Extended Data Figure 4-3.**
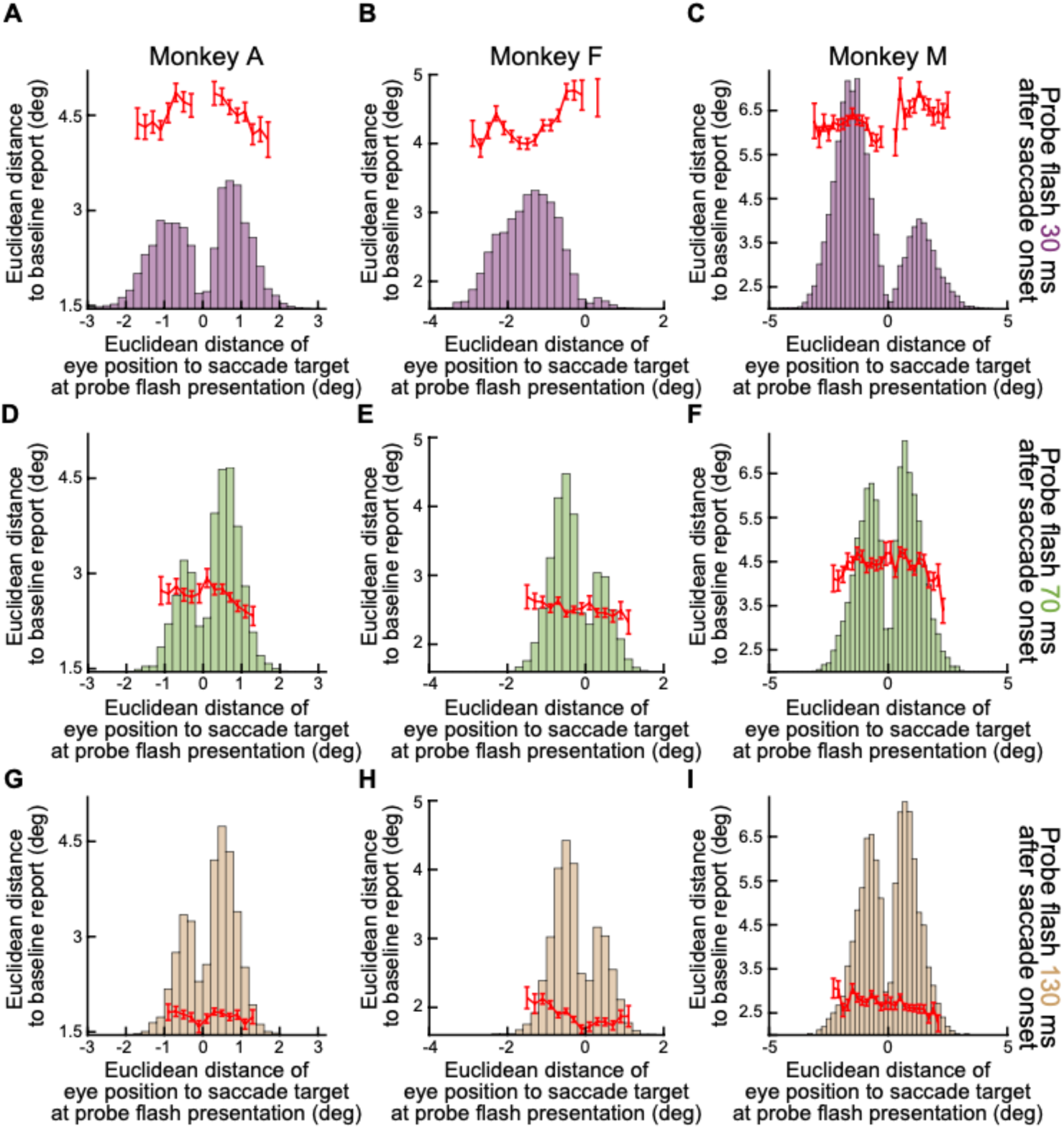
Eye position at the time of probe flash presentation and its relationship to mislocalization strength. **(A-C)** Euclidean distance (red) to the baseline report as a function of eye position at the time of probe flash presentation for probe flashes occurring 30 ms after saccade onset for the individual monkeys. **(D-F)** The same relationship for probe flashes occurring 70 ms after saccade onset. **(G-I)** The same analysis for probe flashes occurring 130 ms after saccade onset. In each case, the faint histograms in the background show the bins in which we had eye position data to document. Across all three monkeys and all time points, no systematic relationship was found between eye position and mislocalization strength, confirming that localization errors were primarily driven by the timing of the flash rather than by momentary gaze position. Indeed, for each monkey, the mislocalization strength decreased much more strongly from the top to bottom row than within each row as a function of eye position.

**Extended Data Figure 4-4.**
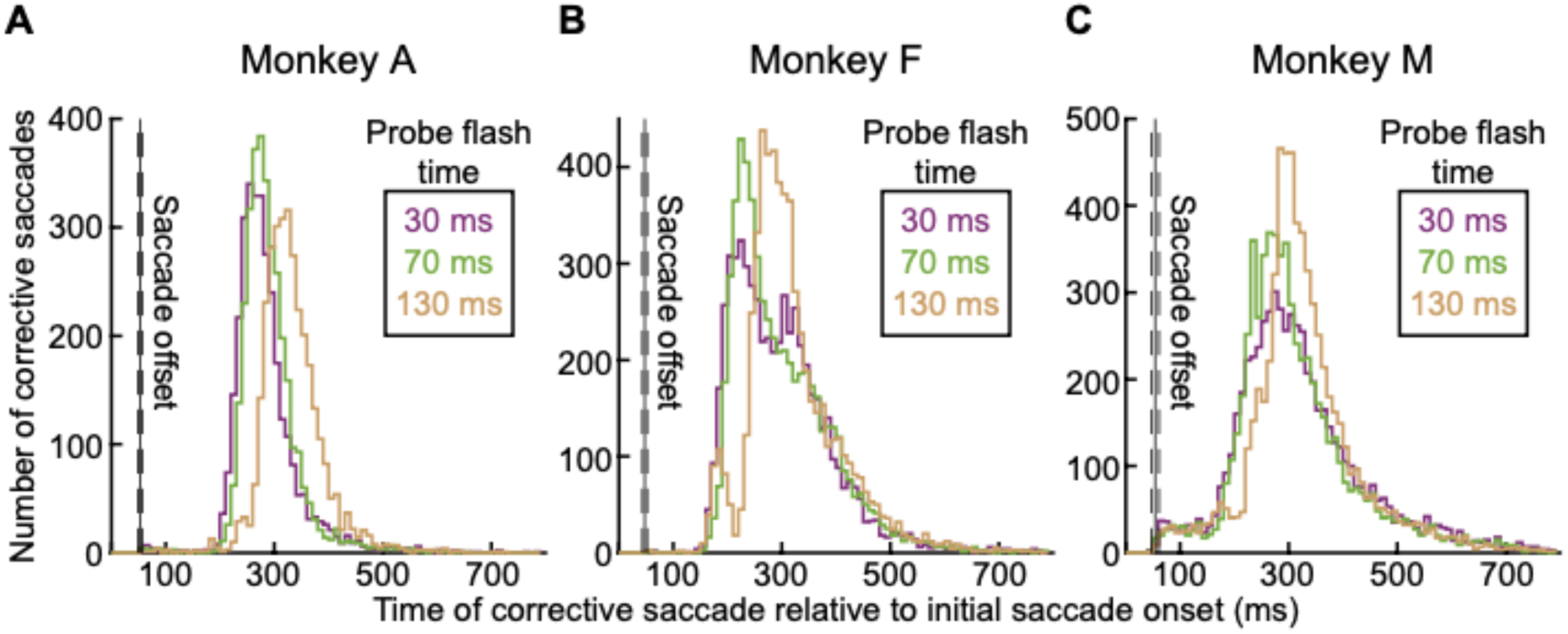
Corrective saccades were too late to account for mislocalization effects. **(A-C)** The temporal distribution of corrective saccades relative to the initial saccade onset for each monkey. The number of corrective saccades is plotted over time, aligned to the initial saccade onset. The different probe flash timings (30 ms, 70 ms, and 130 ms after saccade onset) are color-coded. The majority of corrective saccades occurred after probe flash presentation, confirming that they did not influence perisaccadic mislocalization. A slight delay in corrective saccades for late probe flash presentations (130 ms) suggests transient saccadic inhibition, supporting the notion that the monkeys attended to the probe flash. The vertical black lines with surrounding dashed lines indicate the mean and SEM times of saccade offset.

### Perisaccadic mislocalization was strongest for upward saccades

Having convinced ourselves that we were indeed measuring genuine perisaccadic mislocalization, we were now in a position to explore further properties.

We first focused on the influence of saccade direction. In previous work, we found that backward mislocalization in humans was particularly strong for upward saccades (Grujic et al., 2018). If our current paradigm is robust, and if macaques are indeed a suitable animal model for exploring this property of perisaccadic mislocalization with neurophysiology, then we should also obtain stronger perisaccadic visual mislocalization in our animals for upward saccades. This was indeed the case.

Consider the results of Fig. 5A, B, which are formatted similarly to Fig. 3D. In Fig. 5A, the animal (monkey A) generated purely upward saccades. The mislocalization was backward relative to the saccade direction (also see Fig. 4), and it was stronger for smaller perisaccadic probe flash times than for later times, as expected. Importantly, the mislocalization seen in Fig. 5A was substantially stronger than that observed for purely downward saccades in the same animal. Specifically, in Fig. 5B, we plotted the data for purely downward saccades in the same format as in Fig. 5A. There was still reverse mislocalization opposite the saccade direction, but the peak mislocalization strength (for 30 ms perisaccadic probe flash times) was significantly weaker than in Fig. 5A. This fact is rendered clearer by noting Fig. 5C. Here, we plotted the average (and SEM) reported locations for 30 ms perisaccadic probe flashes for the two saccade directions together in the same plot. To facilitate comparison of the effect sizes in the two cases, we rotated all data to a schematic representation in which the saccade vector was to the right, and the probe flashes were further ahead of it. We used exactly the same approach to visualize results from different saccade directions in our earlier human studies (Grujic et al., 2018). As can be seen, relative to the baseline reports of each condition (black symbols and colored connecting arrows) there was clearly stronger perisaccadic mislocalization (in the backward, reverse direction) for upward saccades. Quantitatively, for purely upward saccades, the average Euclidean distance measure across all three flash locations in this figure was 5.8 deg, whereas it was only 3.5 deg for purely downward saccades. Therefore, there was stronger mislocalization for the upward saccades.

**Figure 5.**
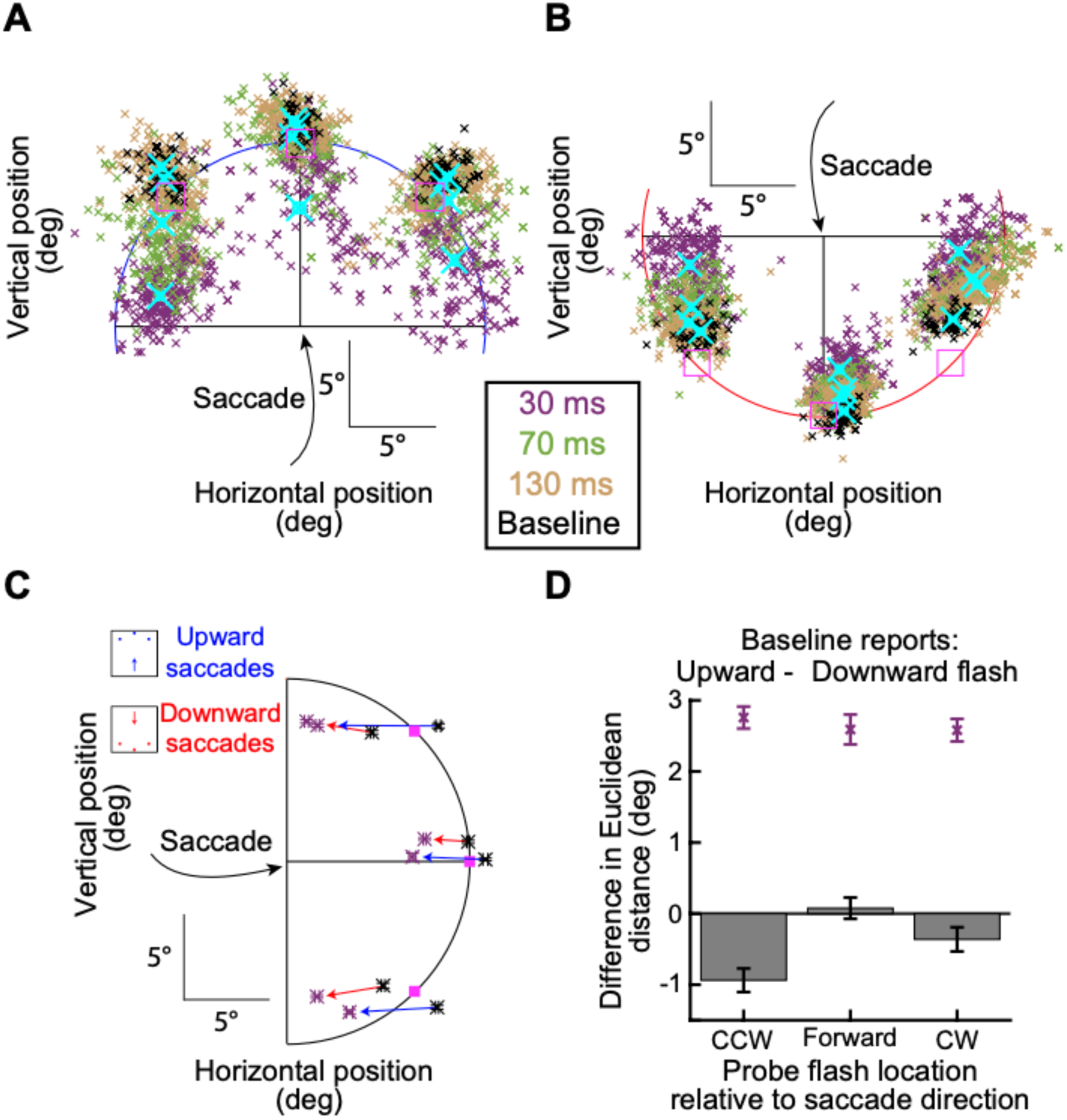
Stronger perisaccadic mislocalization for upward saccades. **(A)** Similar analysis to that in Fig. 3D but for purely upward saccades in monkey A. Reverse mislocalization can be seen, consistent with Fig. 3. **(B)** In the same animal, the strength of reverse mislocalization was weaker for purely downward saccades. This can be seen by the smaller (than in **A**) deviation in the reports for the 30 ms perisaccadic flashes from the baseline reports. Note that we showed the true flash location in **A**, **B** using outline pink squares in this case (unlike in Fig. 3), in order to avoid covering the data points. Otherwise, the figure is formatted like in Fig. 3. **(C)** Comparison of the strength of mislocalization for the two saccade directions in **A**, **B**. Here, we rotated the saccade vectors of **A**, **B** to a single schematic representation (shown here as a rightward saccade), like we did previously (Grujic 0et al., 2018). Purple symbols (along with SEM error bars) indicate reports with the 30 ms perisaccadic flashes, and black symbols (along with SEM error bars) indicate baseline reports. Pink squares indicate the true flash positions. Red and blue connecting arrows indicate whether the saccade was upward (blue) or downward (red). The blue arrows were always longer than the red arrows, indicating a stronger mislocalization (relative to baseline) for the upward than for the downward saccades. **(D)** Baseline reports exhibited minimal differences between upward and downward saccades for each flash location, highlighting that the stronger mislocalization for upward saccades relative to baseline reports (purple points) was not driven by differences in the baseline reports. For the individual probe flash locations, the difference in baseline reports between upward and downward saccades is shown as bars, and the difference in reports at peak mislocalization as purple data points. All error bars denote SEM.

Moreover, this difference in mislocalization strength between upward and downward saccades at peak mislocalization (30 ms flash times) was clearly larger than any systematic differences in the baseline report locations between upward and downward saccades: the Euclidean distance between the baseline report locations (black symbols) for purely upward and purely downward saccades in Fig. 5C was only 0.51 deg (again averaging the respective distances across all three flash locations); this was smaller than the difference in mislocalization strengths between the upward and downward saccades. This result can be seen in Fig. 5D, in which we plotted the difference in baseline reports for upward and downward saccades (for the individual flash locations) as bars; the difference at peak mislocalization between upward and downward saccades is shown as purple data points, confirming that the larger difference in mislocalization strengths between upward and downward saccades was not a consequence of equally large differences in baseline reports. Thus, there was indeed stronger perisaccadic visual mislocalization for upward saccades, replicating previous human observations (Grujic et al., 2018).

We next summarized the above observations across all three animals. In Fig. 6A, B, we measured the Euclidean distance of flash localizations (Fig. 3E, right and Materials and Methods) as a function of probe flash time from saccade onset. We did so separately for all upward saccades and for all downward saccades. In the two shown animals in these two panels, all upward saccades meant purely upward and two oblique upward saccade directions, and all downward saccades meant purely downward and two oblique downward saccade directions (see the inset schematics in Fig. 6A, B). We also included all flash locations per saccade direction. That is, for each saccade direction and saccade location, we collected the Euclidean distance, and we then averaged across all measurements. In both animals, at peak mislocalization (flashes occurring 30 ms after saccade onset in our experiments), there was stronger perisaccadic visual mislocalization for upward saccades, consistent with human observations (Grujic et al., 2018). We also reached the same conclusions when treating each saccade direction individually (Fig. 6C, D).

**Figure 6.**
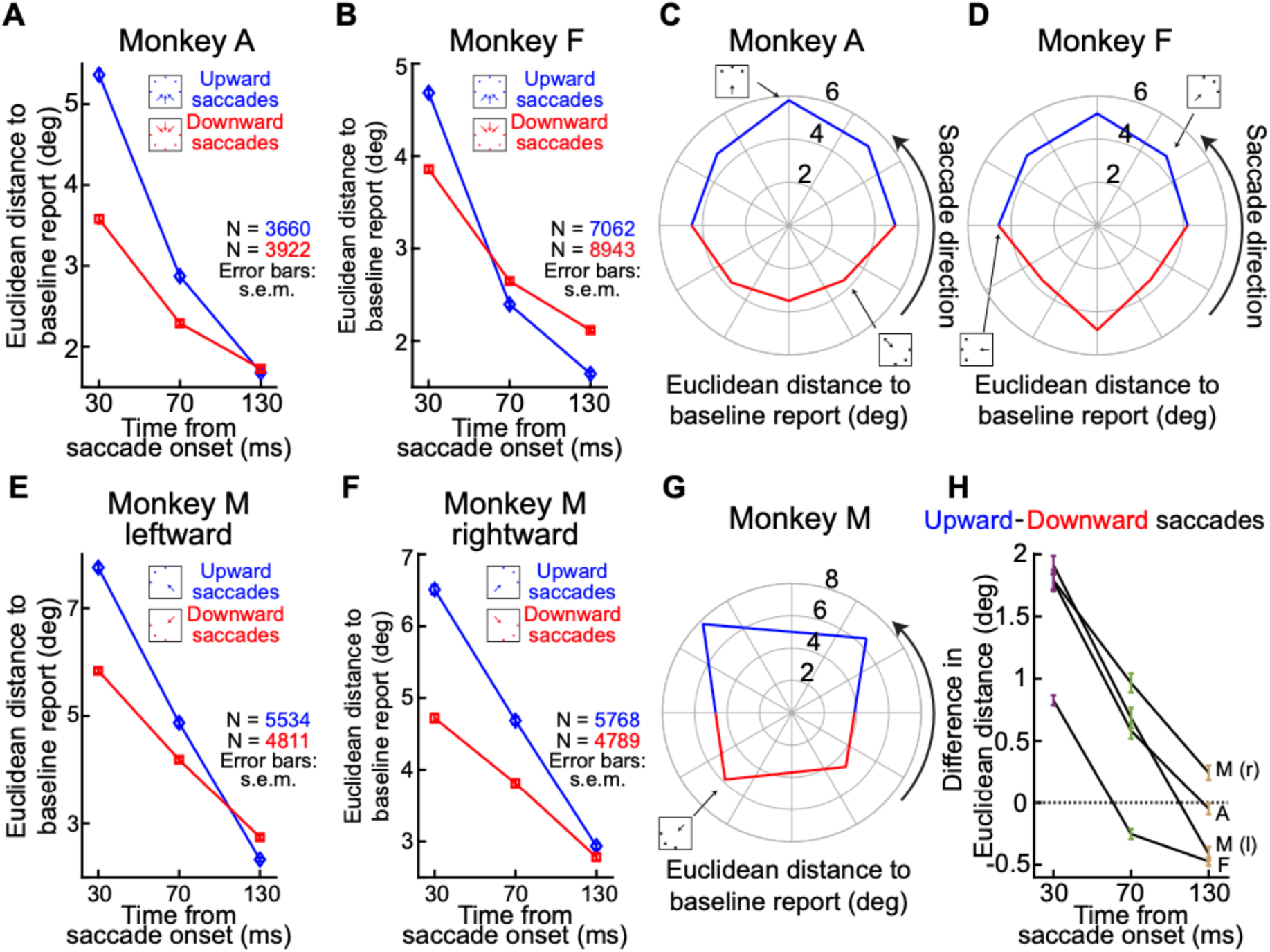
Consistently stronger perisaccadic mislocalization for upward saccades. **(A)** In monkey A, we measured the Euclidean distance of perisaccadic reports (relative to baseline reports) as a function of flash time from saccade onset. The two curves show mislocalization strengths when considering only trials with either upward (blue) or downward (red) saccades. The inset icons clarify the saccade vectors included for each curve (Materials and Methods), as well as their relevant flash locations (3 per saccade direction). There was stronger mislocalization for upward saccades. **(B)** Similar results for monkey F. This monkey also showed more rapid recovery dynamics, with time, for the upward saccades. **(C, D)** Same data as in **A, B,** but considering each saccade direction individually. In each plot, the angle indicates the direction of the saccade (see inset icon examples), and the radial value indicates the Euclidean distance measured for 30 ms flash times. There was stronger mislocalization for upward saccades. **(E, F)** Similar analyses to **A, B,** but for monkey M. In this monkey, we separated saccades with a leftward **(E)** or rightward **(F)** component. In both panels, the monkey still exhibited stronger mislocalization for upward saccades. **(G)** Same as **C, D,** but for monkey M. Again, there was stronger mislocalization for upward saccades. Note that there was also stronger mislocalization for leftward saccades. These were the faster eye movements (approximately 550 deg/s versus approximately 350 deg/s peak velocity), suggesting an interaction between saccade speed and mislocalization strength (Ostendorf et al., 2007). **(H)** We summarized the effects of saccade direction by calculating the difference in mislocalization strength between upward and downward saccades. For all monkeys, there was stronger mislocalization for upward saccades, especially at peak mislocalization times. Note that for monkey M, we show two curves for either leftward (l) or rightward (r) saccades. Error bars indicate SEM.

We also replicated these results in monkey M. In this monkey, we separated the visual hemifields towards which the primary saccades were directed because this monkey had significantly slower saccades with a rightward component, making it difficult to match kinematics across hemifields (Materials and Methods). The monkey again exhibited stronger perisaccadic visual mislocalization for upward saccades (Fig. 6E, F). This can also be seen in Fig. 6G, with the added observation being that perisaccadic visual mislocalization was stronger for the faster leftward saccades. This is also consistent with human observations (Ostendorf et al., 2007).

Statistically, all of the above results were robust at the individual monkey level. We conducted a two-way ANOVA to examine the effects of saccade direction and probe time relative to saccade onset on the Euclidean distance to the baseline report. There was a significant main effect of saccade direction for monkeys A (F(1,7576)=421.07, p<0.001) and M (F(1,20896)=812.77, p<0.001), indicating that the direction of the saccade had a significant impact on perisaccadic mislocalization. Monkey F did not show a significant main effect for saccade direction (F(1,15999)=1.92, p=0.16), but this was only because this monkey exhibited faster recovery times than the other two monkeys (Fig. 6B). Indeed, this monkey, as well as the other two, showed a significant interaction between saccade direction and probe flash time relative to saccade onset (monkey A: F(2,7576)=202.37, p<0.001; monkey F: F(2,15999)=265.66, p<0.001; monkey M: F(2,20896)=333.72, p<0.001), indicating that the effect of saccade direction depended on the time point. There was also a significant main effect of probe time relative to saccade onset in all three monkeys (monkey A: F(2,7576)=1870.19, p<0.001; monkey F: F(2,15999)=3371.03, p<0.001; monkey M: F(2,20896)=4434.12, p<0.001), suggesting that saccadic mislocalization varied across different time points (as expected). Post-hoc comparisons using Bonferroni corrections for probe time relative to saccade onset revealed significant differences between all time points for all monkeys, with visual mislocalization decreasing progressively from 30 to 130 ms (p<0.001). All of these statistical results are summarized graphically in Fig. 6H.

We also quantified mislocalizations for probe flash presentations at 30 ms and 70 ms after saccade onset by measuring the Euclidean distance of these reports not to the baseline trials, but to the reported locations of trials with probe flash presentations at 130 ms after saccade onset. For Monkey A, the average mislocalization error for upward saccades was 4.9 deg ± 0.06 SEM at 30 ms and 2.4 deg ± 0.04 SEM at 70 ms, while for downward saccades, the error was only 3.1 deg ± 0.04 SEM (30 ms) and 1.7 deg ± 0.03 SEM (70 ms). Monkey F (upward saccade: 3.8 deg ± 0.03 deg (30ms), 1.8 deg ± 0.02 deg (70ms); downward saccade: 2.7 deg ± 0.03 deg (30 ms), 1.6 deg ± 0.02 deg (70ms)) and Monkey M (upward saccade(to the left): 6.6 deg ± 0.05 deg (30 ms), 3.8 deg ± 0.05 deg (70 ms); downward saccade (to the left): 4.5 deg ± 0.04 deg (30 ms), 3 deg ± 0.04 deg (70 ms); upward saccade (to the right): 5.6 deg ± 0.05 deg (30 ms), 3.4 deg ± 0.04 deg (70 ms); downward saccade (to the right): 4.3 deg ± 0.05 deg (30ms), 3.1 deg ± 0.04 deg (70 ms)) showed a similar pattern.

Returning to our more robust measures relative to baseline reports, and like we did in Fig. 5D, we also checked the differences in systematic errors in baseline reports themselves. Specifically, since memory-guided saccades of different directions can have different systematic error magnitudes (Hafed and Chen, 2016), we asked whether the differences in mislocalization strengths between upward and downward saccades that we saw in Fig. 6 were entirely due to differences in baseline report errors. For each monkey, we measured the error (from true flash location) in the baseline report saccade as a function of saccade direction (Fig. 7A). We did this only for the baseline trials in which perception was expected to be veridical (Materials and Methods). As anticipated (e.g. Figs. 3, 5), there were non-zero absolute errors in the baseline saccades. However, these errors were significantly smaller than the peak mislocalization effects (also seen in Figs. 3, 5). For example, monkey F had a peak mislocalization amplitude of around 4-5 deg (Fig. 6B), but only a baseline report error of around 1.5-1.6 deg. Most importantly, in all animals, the difference in baseline report errors between upward and downward saccades (Fig. 7B), although significant (monkey A: t(647)=−6.04, p<0.001; monkey F: t(1323)=−5.03,p<0.001; and monkey M: t(689)=−4.11,p<0.001 for leftward saccades and t(704)=−5.78,p<0.001 for rightward saccades; independent samples t-tests), was of the opposite sign from the mislocalization effects comparing upward and downward saccades (purple samples in Fig. 7B and Fig. 6). Thus, stronger perisaccadic mislocalization for upward saccades was not explained by systematic differences in baseline report errors.

**Figure 7.**
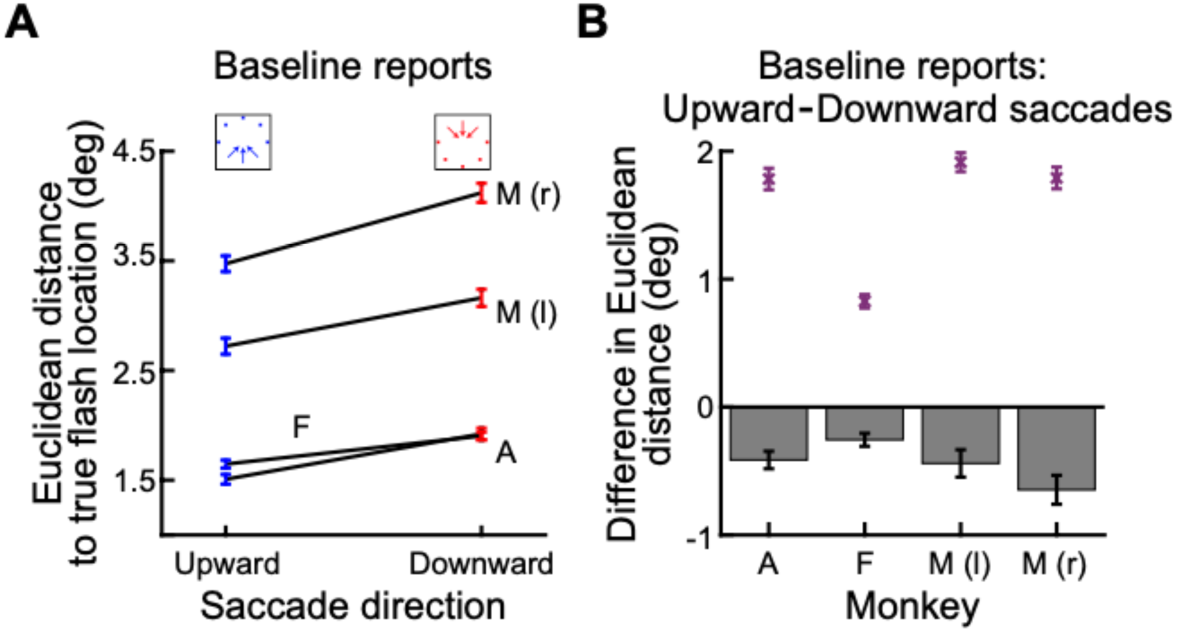
Different patterns of baseline report errors between upward and downward saccades than for mislocalization. **(A)** For each monkey, we measured the Euclidean distance between the baseline report saccade endpoints and the true flash locations in the data of Fig. 6. We did this separately for either upward or downward saccades. Error bars denote SEM. Note how there was still some error from the true flash location in the baseline reports in which no mislocalization was to be anticipated (also visible in Figs. 3, 5). This is expected from memory-guided saccades (White et al., 1994; Willeke et al., 2019; Willeke et al., 2022). However, such error was always smaller than peak mislocalization error (also visible in Figs. 3, 5). **(B)** For each monkey, we took the difference in the errors in **A** between upward and downward saccades (bar plots; similar to Fig. 5D). This difference revealed slightly larger errors for downward memory-guided saccades, consistent with earlier observations (Hafed and Chen, 2016). In contrast, the purple data points are the results from the time of peak mislocalization in Fig. 6. There was a clearly larger difference between upward and downward saccades during peak perisaccadic visual mislocalization than during baseline reports (and also with an opposite sign). Therefore, the results of Fig. 6 were not driven by differences in baseline report errors between upward and downward saccades.

Finally, mislocalization is known to exhibit a two-dimensional landscape, with a component parallel to the saccade direction and another orthogonal to it (see Fig. 8A). In Fig. 8, stronger mislocalization for upward saccades was evident in both directional components. Both the larger mislocalization effect along the saccade direction (Fig. 8B-E) and the smaller localization error orthogonal to the saccade direction (Fig. 8F-I) varied depending on the saccade direction in all three tested monkeys.

**Figure 8.**
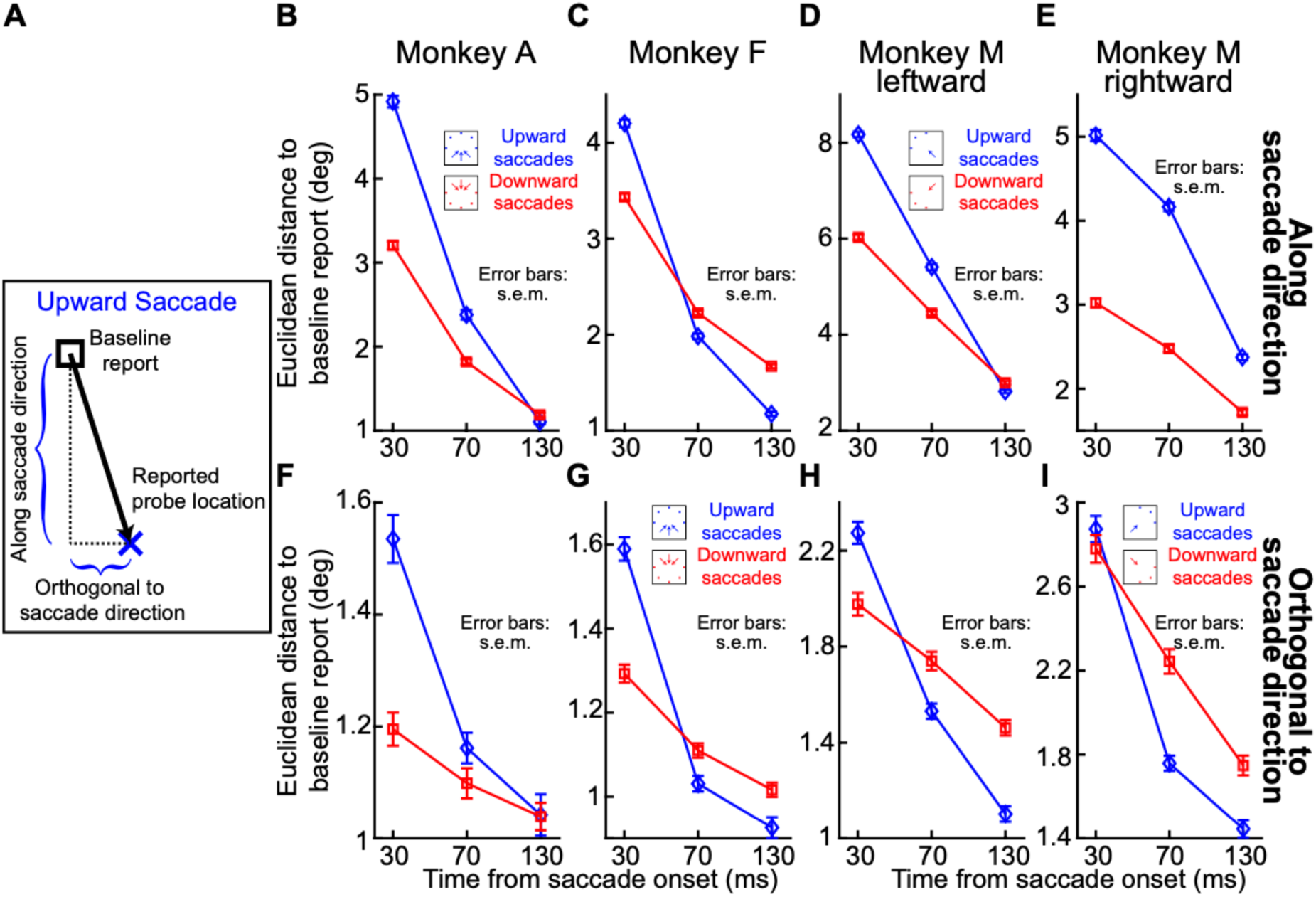
Two-dimensional perisaccadic mislocalization landscape in macaques. **(A)** our analysis approach for assessing the strength of orthogonal mislocalization (Baumann et al., 2024). We measured the two components of mislocalization separately. One component was along the saccade direction, and one component was orthogonal to the saccade direction (Baumann et al., 2024). **(B-E)** Mislocalization along the saccade direction for upward (blue) and downward (red) saccades in the three monkeys (A, F, M). Note that we show rightward and leftward saccade directions separately for Monkey M. Error bars represent SEM. There was consistently stronger mislocalization for upward saccades. **(F-I)** Localization error orthogonal to the saccade direction for the same conditions. Stronger mislocalization for upward saccades was still evident, with a pronounced effect along the saccade direction and a smaller but consistent error orthogonal to the saccade direction across all tested monkeys.

Therefore, our results so far indicate that all three monkeys experienced perisaccadic visual mislocalization, and that they all demonstrated an effect of saccade direction, consistent with human observations (Grujic et al., 2018). A quantitative comparison of effect magnitudes as a function of saccade speeds was also afforded to us by monkey M, and the results were again consistent with human observations (Ostendorf et al., 2007). This reinforces our interpretation of the flexibility and robustness of our paradigm.

### Perisaccadic mislocalization was stronger for upper visual field flashes

We next explored the effect of visual field location of the flashes relative to horizontal saccades. In humans, we recently found that, with such horizontal saccades, reverse mislocalization was stronger for upper visual field flashes (ahead of the saccade target location) than for symmetrically located lower visual field ones (Baumann et al., 2024). This was also the case in our monkeys. Specifically, in Fig. 9A-D, we analyzed mislocalization strength for only horizontal saccades in each monkey. This time, we separated the trials based on whether the perisaccadic flash location was above or below the horizon. That is, if the saccade was to the right, our paradigm included three potential flashes: one directly ahead of the saccade target (still on the horizontal axis), and two oblique ones (Fig. 1, Materials and Methods). The two oblique flashes were, by definition, either in the upper or lower visual field relative to the saccade direction, and we compared these two oblique flash positions in the current analysis. The results demonstrated that there was indeed generally stronger mislocalization associated with upper rather than lower visual field flashes.

**Figure 9.**
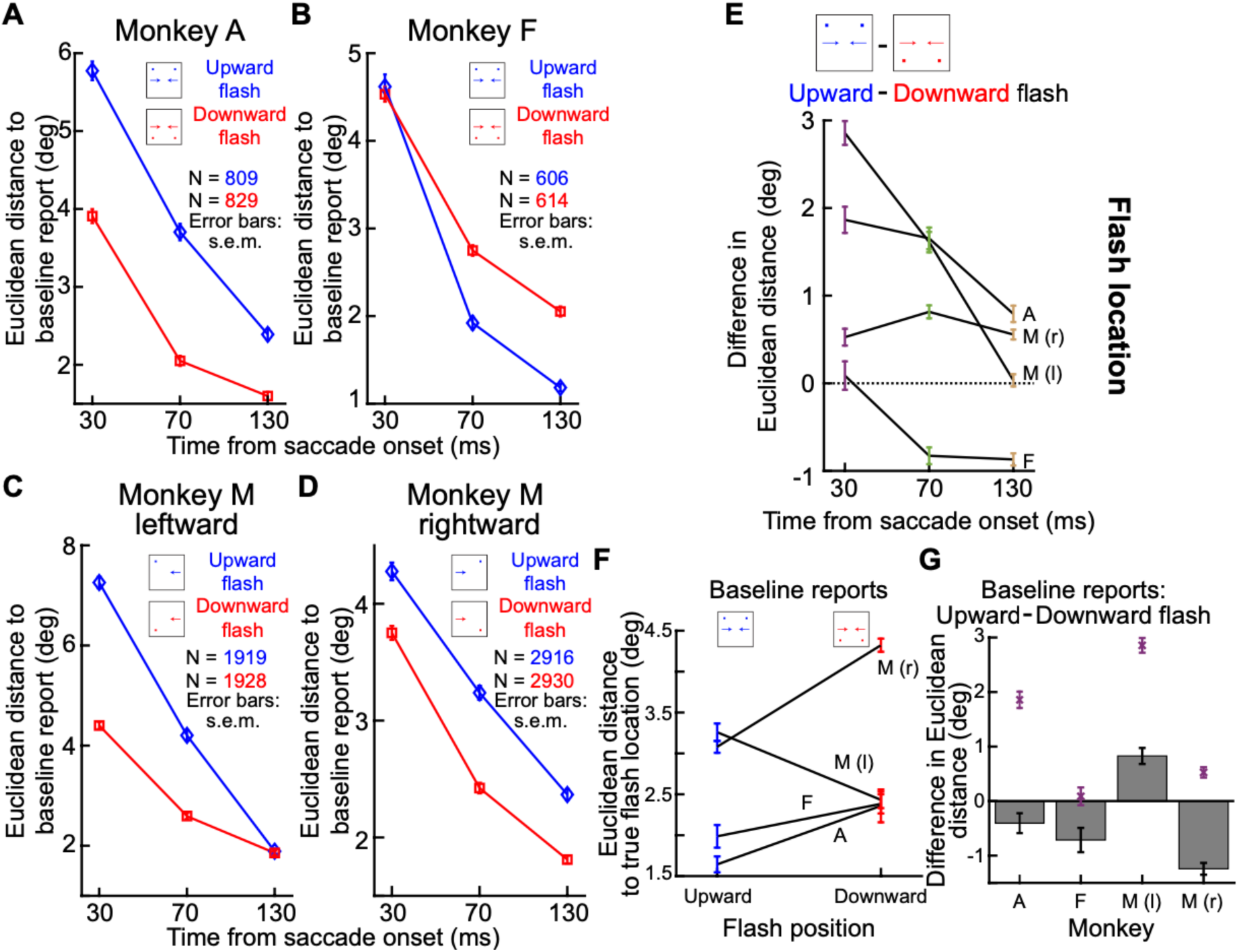
Stronger perisaccadic visual mislocalization for upper visual field probe flashes and horizontal saccades. **(A)** For monkey A, we only considered horizontal primary saccade directions. We then compared mislocalization strength (Euclidean distance to baseline reports) for probe flashes above or below the saccadic vector (see Materials and Methods and inset schematic). There was stronger mislocalization for upper visual field flashes, like in humans (Baumann et al., 2024). **(B)** In monkey F, the effect at peak mislocalization (flashes 30 ms from saccade onset) was much weaker than in monkey A, but with a similar trend. This monkey also had rapid time dynamics, like in Fig. 6, causing a reversal at longer times. **(C, D)** Monkey M replicated the results of monkey A regardless of the direction of the horizontal saccades. **(E)** Summary of the results in **A**-**D** shown here as the difference in mislocalization strength between upper and lower visual field flash locations, and as a function of flash time (like in Fig. 6H). **(F)** Errors in baseline reports from true flash location, like in Fig. 7A. Here, we plotted these errors for either the upward or downward flash conditions. The errors were always smaller in magnitude than the peak mislocalization effects in **A**-**E** (consistent with Figs. 3, 5). **(G)** More importantly, the difference in these errors between upward and downward flashes did not explain the differences in peak mislocalization. This panel is formatted similarly to Fig. 7B: the bars denote the differences in baseline report errors between upper and lower visual field flashes, and the purple symbols show the peak mislocalization differences from **A**-**E**. Differences in baseline reports did not explain differences in peak mislocalization strengths (except for monkey F where there were already no differences in mislocalization strength). Error bars denote SEM.

Statistically, we performed a two-way ANOVA to evaluate the effects of probe location and probe time relative to horizontal saccade onset on the Euclidean distance to the baseline report. There was a significant main effect of probe location for all three monkeys (monkey A: F(1,1632)=410.01, p<0.001; monkey F: F(1,1214)=65.2, p<0.001; monkey M: F(1,9687)=626.66, p<0.001) indicating that the probe location had an impact on the strength of perisaccadic visual mislocalization. The main effect of probe time relative to saccade onset was also significant (monkey A: F(2,1632)=552.54, p<0.001; monkey F: F(2,1214)=714.44, p<0.001; monkey M: F(2,9687)=1652.95, p<0.001), suggesting that perisaccadic mislocalization varied across different time points. Finally, the interaction between probe location and probe time relative to saccade onset was also significant for all three monkeys (monkey A: F(2,1632)=21.43, p<0.001; monkey F: F(2,1214)=21.93, p<0.001; monkey M: F(2,9687)=70.51, p<0.001), indicating that the effect of probe location was dependent on flash time. These results are summarized in Fig. 9E showing difference measures similar to ones that we used earlier for summarizing saccade direction effects.

We also checked whether systematic differences (between upper and lower visual field flashes) in baseline report errors accounted for the above results. Specifically, just like in Fig. 7A, we plotted the Euclidean distance between baseline reports and true flash location for the upward and downward flash conditions of this analysis. The results are shown in Fig. 9F. Consistent with Fig. 7A, the absolute errors in baseline reports were smaller than the peak mislocalization effects; this is also consistent with the raw example data shown in Figs. 3, 5, demonstrating clear recovery with time from saccade onset (towards the baseline report locations). Importantly, the differences in baseline report errors between upward and downward flashes were not correlated with the differences in peak mislocalization strength (Fig. 9G). Thus, the effects of upper and lower visual field flashes in association with horizontal saccades were likely perisaccadic and not an artifact of report saccade errors.

### Perisaccadic mislocalization strength was subtly modulated by the saccade target appearance

Finally, we explored whether mislocalization strength depended on the visual appearance of the saccade target (Materials and Methods). In two out of the three monkeys, there was a very subtle difference in mislocalization strength as a function of saccade target appearance; however, this dependence was not the same in both animals. Consider for example, Fig. 10A. The figure shows slightly stronger mislocalization for the low spatial frequency grating as the saccade target. However, Monkey F showed the opposite effect (Fig. 10B). Figure 10C, D shows the data of monkey M, showing no image dependency for the time bin with peak mislocalization, and neither for leftward nor for rightward saccades. These findings for the image dependence are smaller than the dependence for saccade direction.

**Figure 10.**
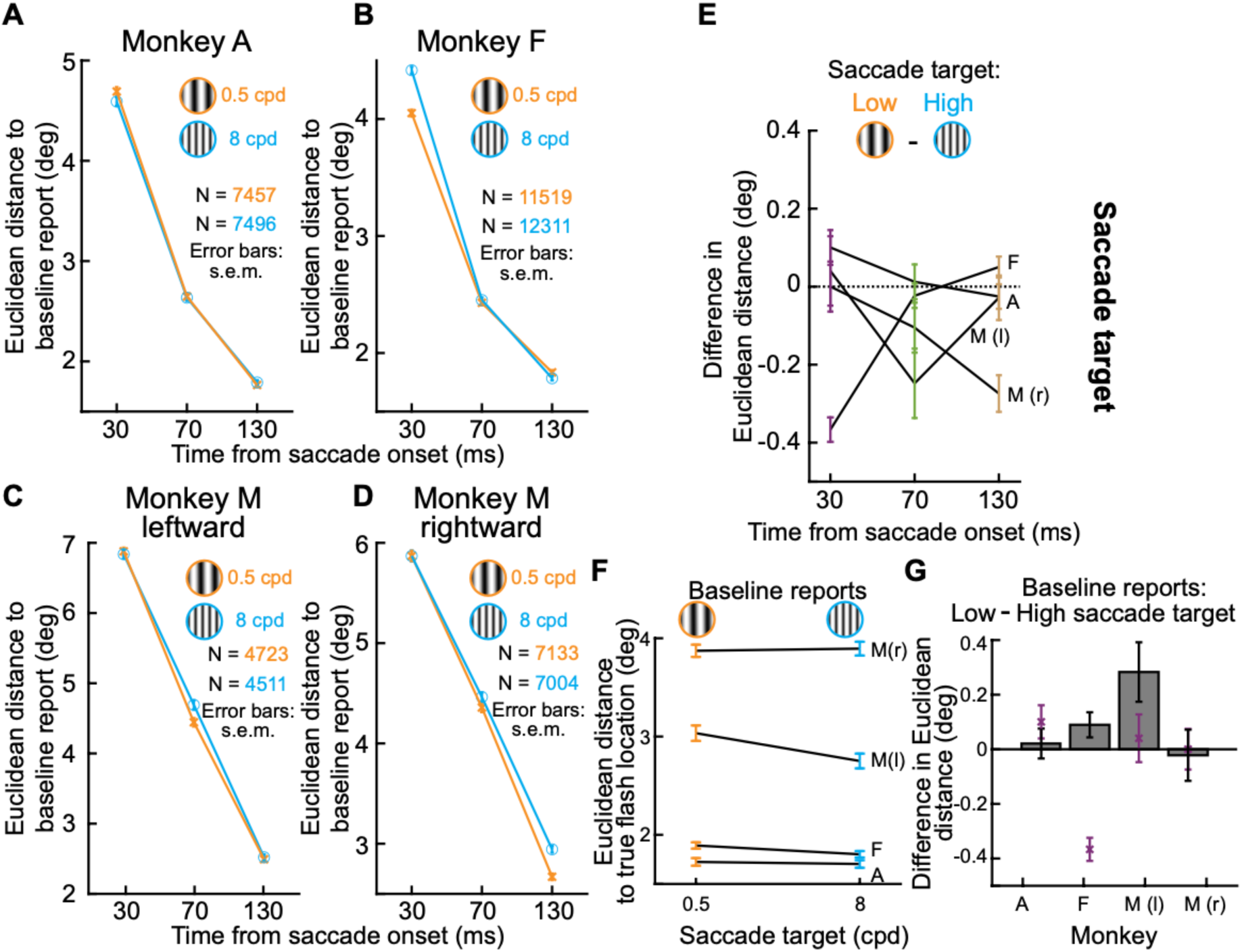
Inconsistent effects with respect to saccade target appearance. This figure is formatted identically to Fig. 9. Here, the comparison was between the different saccade target appearances. Therefore, we pooled across saccades of all directions (Materials and Methods) and only separated trials based on the appearance of the saccade target. There were subtle differences **(A-E)**, but the differences were similar to the differences in baseline report saccades in the absence of perceptual mislocalization **(F, G)**. Thus, the image dependence effect in perisaccadic mislocalization (Baumann et al., 2024) was likely masked by systematic errors in report saccades. All other conventions are similar to Fig. 9.

Statistically, we performed a two-way ANOVA to investigate the effects of saccade target appearance and probe time relative to saccade onset on the Euclidean distance to the baseline report. There was a significant main effect of saccade target appearance for two of the three monkeys (monkey A: F(1,14947)=1.15, p=0.28); monkey F: F(1,23824)=33.05, p<0.001; monkey M (F(1,21454)=20.32, p<0.001) indicating that the saccade target had an impact on the strength of perisaccadic mislocalization. However, monkeys’ M and F results were not consistent with the previous human results: these monkeys exhibited stronger mislocalization for a high spatial frequency grating. As expected, there was also a significant main effect of probe time relative to saccade onset (monkey A: F(2,14947)=3867.51, p<0.001; monkey F: F(2,23824)=5413.83, p<0.001; monkey M: F(2,21454)=4889.48, p<0.001), suggesting that saccadic mislocalization varied across different time points. Additionally, the interaction between saccade target appearance and probe time relative to saccade onset was significant for all three monkeys (monkey A: F(2,14947)=1.87, p<0.001; monkey F: F(2,23824)=42.36, p<0.001; monkey M: F(2,21454)=4.77, p<0.001).

Thus, we cannot confidently conclude that there was an image dependence of mislocalization strength in our monkeys. This is likely caused, at least in part, by the differences in the ways monkeys and humans reported their percepts (see Discussion). Indeed, analyzing the baseline report errors was particularly revealing (Fig. 10F, G): the difference in magnitudes of baseline report errors between the different target image appearances (Fig. 10G) was of the same order of magnitude as the difference in the peak mislocalization strengths for the different saccade target images. This suggests that a possible image dependence of perisaccadic mislocalization in our monkeys might have been masked by errors in the memory-guided report saccades.

## Discussion

We established a robust paradigm with which we could probe perisaccadic mislocalization (with clear visual references in the environment) in monkeys. Importantly, our paradigm allows mapping the two-dimensional landscape of mislocalization, and it can also be used with arbitrary saccade directions. This latter property enabled us to replicate the observation in humans (Grujic et al., 2018) that backward perisaccadic “compression” of flashes ahead of the saccade target is strongest for upward saccades. This dependency of mislocalization on saccade direction was also visible in both directional components, along and orthogonal to the saccade direction. Moreover, we replicated the human observation (Baumann et al., 2024) of stronger backward mislocalization for upper visual field stimuli during horizontal saccades.

Our results can be compared to human psychophysics conducted under similar visual reference conditions. Specifically, a biphasic perisaccadic mislocalization pattern, as reported by (Honda, 1991, 1993), is a well-established phenomenon observed primarily under complete darkness. In this case, when flashes are presented immediately before or at the onset of a saccade, mislocalization typically occurs in the same direction as the saccade. Conversely, when flashes are presented at the end or immediately after the saccade, the perceived location of the flash shifts in the direction opposite to the saccade direction, resulting in a characteristic forward and then backward pattern of mislocalization. However, the presence of visual references appears to significantly modulate this effect. For example, (Honda, 1999) demonstrated that introducing even minimal visual context, such as a faint frame pattern or a dimly illuminated structured background, substantially reduced mislocalization and disrupted the classic biphasic pattern. Under these conditions, mislocalization was more akin to a “compression” effect, where the perceptual error depended on the flash position relative to the saccade target rather than purely on the temporal relationship to the saccade onset. These findings align with the results of studies by (Ross et al., 1997) and (Lappe et al., 2000), which also showed that structured visual environments promoted “compression”-like mislocalization patterns. Since we included a stable visual reference (the saccade target) and bright display conditions, our results are consistent with this framework. Of course, in future work, we can confirm this explicitly by testing probe flash locations in between the initial and final saccade fixation points (Grujic et al., 2018).

Our work builds on previous demonstrations that perisaccadic visual mislocalization can be studied in macaques (Dassonville et al., 1992; Jeffries et al., 2007; Klingenhoefer and Krekelberg, 2017; Weng et al., 2024). Our work also relates to interesting monkey variants of the classic double-step saccade task (Hallett and Lightstone, 1976; Becker and Jurgens, 1979; Ottes et al., 1984; Vliegen et al., 2005), in which the target of the second saccade in the double-step sequence was presented during the progression of the first eye movement (and thus could be susceptible to some mislocalization) (Van Grootel et al., 2012). However, in the context of visual mislocalization studies, our approach adds paradigm flexibility that we believe is necessary for further success in porting these behavioral measures of perisaccadic mislocalization to neurophysiology. For example, it may not always be possible neurophysiologically to probe only horizontal flash position mislocalization or only horizontal saccades. This is because electrode positions in topographic maps may be in off-axis representations. More importantly, mislocalization has a two-dimensional landscape (Kaiser and Lappe, 2004). Thus, with our work, we provide an additional stepping stone towards direct neurophysiological experiments exploiting essentially identical behavioral tasks to those classically used in humans.

Indeed, we were motivated by the particularly strong backward mislocalization for upward saccades in humans (Grujic et al., 2018), and that we think deserves further investigation. We suspect that this phenomenon might be related to the asymmetry of representation in the superior colliculus between the upper and lower visual fields (Hafed and Chen, 2016; Zhang et al., 2022; Hafed, 2025), but this needs to be tested explicitly. In fact, besides recording activity related either to the saccade vector endpoint or to flash locations, we could make variants of our behavioral paradigm that can be more causal in nature. For example, we could replace the flash presentation with brief electrical microstimulation of the primary visual cortex, or higher-level cortical areas. Similarly, we could replace the visually-guided saccade with an electrically evoked saccade from the superior colliculus. This latter experiment would be particularly fascinating. For example, it is known that beyond a certain strength of electrical microstimulation in the superior colliculus, evoked saccades become more similar to each other under different stimulation parameters (Katnani and Gandhi, 2012). Thus, one could stimulate with two different “motor burst strengths” and ask how perisaccadic flashes are mislocalized under the two conditions (with the evoked saccades being the same). This would causally establish a functional role for sensory tuning in collicular motor bursts (Baumann et al., 2023) in perisaccadic perceptual phenomena that might, at least theoretically (Baumann et al., 2024), rely on saccade-related corollary discharge.

Similarly, we could perturb brain areas not implicated in corollary discharge and identify the limits of visual contributions to perisaccadic mislocalization. For example, if monkey M’s asymmetry in saccade peak speeds that we alluded to above was due to a region downstream of the superior colliculus, then it could be that the saccade motor commands (from the colliculus), and their associated (colliculus-related) corollary discharge, were the same for either rightward or leftward saccades in this monkey. If so, then the differences that we saw in the magnitude of perisaccadic mislocalization across hemifields in this animal would have been intrinsically visual in origin: the monkey would have had similar saccade commands (at the collicular level) but different speeds of retinal-image shifts. The above example variants of our paradigm demonstrate why we think that this paradigm is useful for neurophysiology, and also why we think that it can meaningfully add to a wealth of previous work exploring perisaccadic modulations in visual processing in the brain (Duhamel et al., 1992; Nakamura and Colby, 2002; Krekelberg et al., 2003; Kusunoki and Goldberg, 2003; Sommer and Wurtz, 2006; Churan et al., 2011, 2012; Zirnsak et al., 2014; Neupane et al., 2016b, a, 2020; Qian et al., 2022). These prior experiments did not exploit a simultaneous behavioral report by the animals, which our paradigm allows.

Of course, and as mentioned above, we do acknowledge that we did not explicitly test flash positions between the initial and final fixation positions associated with the saccade (which should show forward mislocalization). However, our use of multiple different saccade directions, coupled with the consistent backward mislocalization seen in all cases, makes us confident that what we saw was indeed that the prior literature referred to as “compression”. Indeed, the vectors of mislocalization that we observed (e.g. Figs. 3, 5) as a function of time (relative to the saccade vector direction) were similar to those that we observed earlier in humans using multiple different saccade directions (Grujic et al., 2018). Thus, we believe that our results are consistent with those of (Klingenhoefer and Krekelberg, 2017). Importantly, our paradigm can be easily modified to also include flashes nearer to the initial fixation position than the saccade target.

We also agree with (Klingenhoefer and Krekelberg, 2017) that there can (and should) be quantitative differences in the parameters of the observed perisaccadic mislocalization between monkeys and humans. This is totally expected because factors like foveal magnification and sensory transduction speeds, and others, can be quantitatively different between the two species, even when they are qualitatively similar. Such quantitative differences are inevitable given the different brain sizes. What is important is that the qualitative equivalence is present, and this motivates using these animals in particular for invasive neurophysiological investigations.

One caveat with our work is that we relied on memory-guided saccades to report flash location. Such saccades have systematic errors in their landing positions (White et al., 1994; Willeke et al., 2019), which could also depend on the visual field location of the flashes (Hafed and Chen, 2016). Such saccades also have larger dispersions across trials (variable errors) than visually-guided movements. Still, we chose to use memory-guided saccades because we did not want to impose any prior assumptions about where the monkeys were going to see the flashes. For example, if we had used “choice” targets, then we would have been imposing constraints on the spatial reports of the animals. This is problematic for a general two-dimensional paradigm like the one that we were trying to develop. Moreover, even manual pointing (via displayed cursors) by humans has systematic and variable errors (Willeke et al., 2019), and such pointing is generally used in mislocalization experiments (Grujic et al., 2018; Baumann et al., 2024). Thus, some amount of reporting errors is inevitable in any paradigm requiring a behavioral response from the subjects.

We also acknowledge that our choice to use memory-guided saccades could have come at the expense of having enough resolution to observe more subtle effects, like a potential dependence of perisaccadic mislocalization strength on the saccade target image appearance. Indeed, in parallel experiments with humans, we did find more systematic effects across most subjects than with the monkeys in the current study (Baumann et al., 2024). Thus, the inconsistency of the image dependence effects that we saw with our monkeys (Fig. 10) could reflect either added contamination by errors in memory-guided saccades, species differences from humans, or the need to test more monkeys with our current paradigm before reaching a conclusion about an image dependence of perisaccadic mislocalization in the animals.

One possibility to avoid the noise associated with memory-guided saccades could be to modify our paradigm slightly: we could adopt the approach of (Klingenhoefer and Krekelberg, 2017; Weng et al., 2024). That is, at the end of every trial, we would display a very dense two-dimensional array of grid positions, and the monkeys would simply choose which grid position matched the perceived flash location. We do not expect this to cause an increased training burden to the task. In any case, for the effects of saccade directions on mislocalization strengths, our paradigm was sufficient.

We also chose to not test presaccadic flashes. This was motivated purely by the pragmatic decision to increase data yield. By presenting flashes contingent on detecting a saccade, we were essentially ensuring obtaining a usable data point for every saccade trial accepted into the analysis. On the other hand, if we had randomized flash times independent of the saccadic reaction time, we would have needed to collect more trials, and then recalculate flash times relative to saccade onset. This should work since it does not alter the paradigm dramatically, and it can also allow obtaining a full time course for perisaccadic mislocalization in monkeys. Moreover, we have previously used both approaches successfully in our experiments on perisaccadic visual suppression, in both humans and monkeys (Hafed and Krauzlis, 2010; Chen and Hafed, 2017; Idrees et al., 2020), and also in our perisaccadic perceptual mislocalization experiments in humans (Grujic et al., 2018; Baumann et al., 2024). Having said that, one has to carefully interpret trials in which the flash happened too long before saccade onset, especially because such a condition gives rise to a complete resetting of saccade plans (Buonocore and Hafed, 2023).

In all, we established a robust, two-dimensional perisaccadic mislocalization paradigm that is particularly useful for neurophysiological experiments with macaque monkeys.

## Acknowledgements

We thank Tong Zhang, Tatiana Malevich, and Antimo Buonocore for help in data collection.

## Conflict of interest

Authors report no conflict of interest.

## Funding sources

We were funded by the Deutsche Forschungsgemeinschaft (DFG; German Research Foundation) through the SFB 1233 Robust Vision (project number: 276693517) as well as the SPP 2205 Evolutionary Optimization of Neuronal Processing (project number: HA 6749/3-2) and the SPP 2411 Sensing LOOPS: Cortico-subcortical Interactions for Adaptive Sensing (project numbers: 520617944 and 520283985, HA 6749/11-1).

## Notes

### Competing Interest Statement

The authors have declared no competing interest.

### Summary of Updates

Added Figs. 4, 8, as well as extensive control analyses for effects of memory duration, eye position, corrective saccades, and timings.

